# Two D-loop resolution systems enable natural genetic transformation in bacteria

**DOI:** 10.1101/2024.02.06.579203

**Authors:** Léo Hardy, Violette Morales, Clothilde J. Rousseau, Mathieu Bergé, Fanny Mazzamurro, Julie Plantade, Triana N. Dalia, Ankur B. Dalia, Eduardo P.C. Rocha, Xavier Charpentier, Patrice Polard

**Affiliations:** CIRI, Centre International de Recherche en Infectiologie, Inserm, U1111, Université Claude Bernard Lyon 1, CNRS, UMR5308, École Normale Supérieure de Lyon, Univ Lyon, 69100, Villeurbanne, France; Laboratoire de Microbiologie et Génétique Moléculaires, UMR5100, Centre de Biologie Intégrative, Centre Nationale de la Recherche Scientifique, 31062 Toulouse, France; Université de Toulouse, 31062 Toulouse, France; Institut Pasteur, Université Paris Cité, CNRS UMR3525, Microbial Evolutionary Genomics, Paris, 75015, France; Collège Doctoral – Sorbonne Université, F-75005 Paris, France; Department of Biology, Indiana University, Bloomington, IN 47405, USA

## Abstract

Natural transformation is a widespread mechanism driving genetic exchanges in bacteria. It proceeds by the capture and internalization of exogenous DNA in linear single strands, ultimately integrated in the genome by homologous recombination. It is unknown how the RecA-directed D-loop intermediate of this dedicated recombination pathway is processed. We report that resolution of the transformation D-loop depends on two endonucleases of opposing phylogenetic distribution in bacteria. One is YraN, which has co-evolved and interacts with the ComM helicase, known to extend DNA recombination at the transformation D-loop. The other is CoiA, which is restricted to the Bacillota. CoiA is shown to be a resolvase of the transformation D-loop, extended by the RadA helicase in these species. We demonstrate that both YraN and CoiA act synergistically with their cognate helicases. These findings reveal that bacteria have evolved two helicase/nuclease pairs for the maturation and recombination extension of the transformation D-loop.

---

One of the remarkable features of bacteria is their ability to naturally undergo natural genetic transformation, a widespread and conserved mechanism of horizontal gene transfer (HGT) (*1*). Natural transformation is intrinsic to bacteria: it is a chromosomally encoded mode of HGT that, unlike transduction and conjugation, does not rely on the infectious and propagative behaviors of mobile genetic elements. It allows bacteria to import exogenous DNA and integrate it into their chromosome by homologous recombination (Fig. 1A). In many bacteria, the genes involved in DNA uptake and recombination during natural transformation are co-regulated and expressed during a differentiation state known as competence, which is controlled by species-specific regulatory pathways (*1*).

**Fig. 1.**
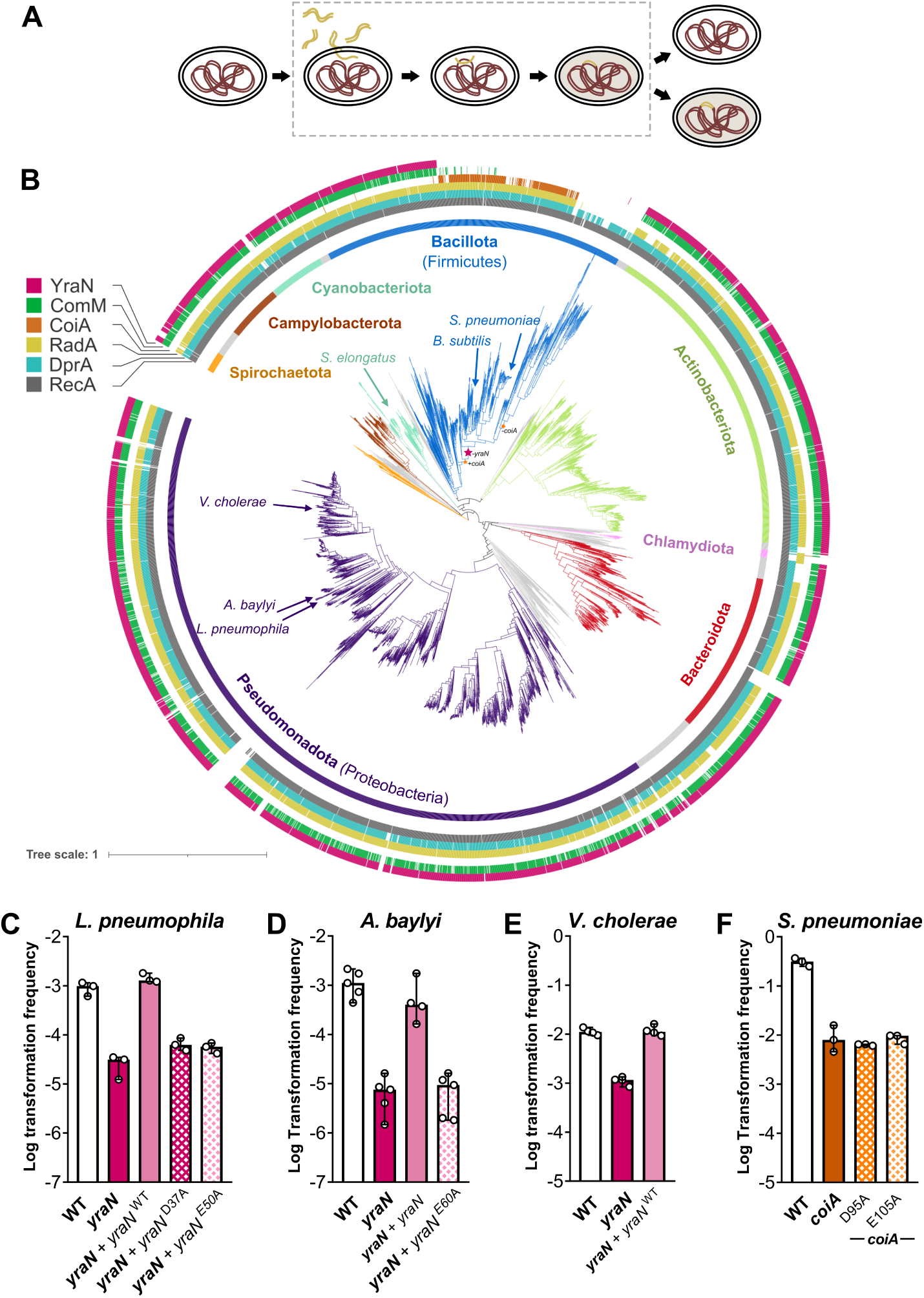
CoiA and YraN are widespread PD-(D/E)XK enzymes involved in natural transformation. (A) Schematic representation of natural transformation. Upon entry into the cytoplasm, the double-stranded tDNA (gold) is converted to ssDNA. A D-loop is formed by tDNA invasion of the chromosome (dark red), creating an heteroduplex intermediate. After resolution of the D-loop, replication and division, one daughter cell inherits the modification conferred by the tDNA while the other daughter cell stays as parental. (B) Distribution of selected transformation-associated proteins in the bacterial phylogenetic tree. The tree consists of 4,185 strains subsampled from the 21,105 bacterial genomes searched for the presence of CoiA, YraN, RadA, ComM, DprA, and RecA. The taxonomy was provided by Genome Taxonomy Database (GTDB). The stars represent *coiA* gain and loss (orange) and *yraN* loss (pink). (C), (D) and (E) Efficiency of natural transformation of a SNP conferring resistance to streptomycin (*L. pneumophila*, *Lp*) or rifampicin (*A. baylyi*, *Ab*, and *Vibrio cholerae*, *Vc*). Parent strain (WT), mutants of *yraN*, and complementation clones were exposed to transforming DNA consisting of a 4 kb-long PCR product encompassing a *rpsL* allele conferring resistance to streptomycin for *Lp*, an 11 kb-long or 5 kb-long PCR product encompassing a *rpoB* allele conferring resistance to rifampicin for *Ab* and *Vc*, respectively. The complementing *yraN* allele in *Lp* (“+*yraN*”) is placed in the endogenous plasmid pLPP and expressed from a constitutive promoter. The complementing *yraN-flag* allele (“+*yraN*”) in *Ab* is placed in the pMMB207C plasmid under control of an IPTG-inducible promoter, tested at 1 mM IPTG (also see Fig. S1D). The complementing *yraN* allele in *Vc* (“+*yraN*”) is placed on a pBAD plasmid and 0.2% arabinose was used for its expression. D37A and E50A correspond to mutations introduced in the catalytic site of YraN of *Lp*. E60A in YraN of *Ab* is same mutation as E50A in YraN of *Lp*. Data represent the median with range of three (*Lp*) and five (*Ab*) independent experiments. (F) Efficiency of natural transformation of a SNP conferring resistance to rifampicin in *S. pneumoniae* (*Sp*). Parent strain (WT), deletion mutant of *coiA*, and mutants of the active site D95A and E105A were exposed to transforming DNA consisting of a 4 kb-long PCR product of the *rpoB* allele in which the mutation conferring resistance is off-centered. Data represent the median with range of three independent experiments.

The DNA processing steps driving its transport across the cell envelope and the cytoplasmic membrane up to its integration in the chromosome are well conserved between monoderms (bacteria with a single cell membrane, like most Gram-positive) and diderms (bacteria with an outer membrane like most Gram-negative) (*2*). The process is initiated by the capture of exogenous double-stranded DNA (dsDNA) by an extracellular Type IV filament (*3*, *4*). While in diderms, the Type IV filament serves other functions such as motility and adhesion (*5*), the competence-associated Type IV filament of monoderms is, to the best of our knowledge, dedicated to the transformation pathway. In both diderms and monoderms, retraction of the filament conveys DNA to the cytoplasmic membrane and is essential for transformation (*3*, *6*, *7*). There, the ComEA DNA receptor functions as a ratchet for dsDNA (*8–10*). The dsDNA is converted into ssDNA as it is transported across the cytoplasmic membrane, presumably through a channel formed by the ComEC protein (*11–13*). In diderms, the ComFC protein is also required for this transport stage, yet the function of this protein remains elusive (*14–17*). Upon entry in the cytoplasm, the ssDNA is rapidly bound by DprA, a transformation-dedicated ssDNA-binding protein widely conserved in bacteria. DprA forms a nucleocomplex protecting the ssDNA from nucleases and mediates RecA polymerization, which initiates a search for sequence homology in the genome (*18*, *19*). If sufficient sequence identity is found, RecA-dependent strand invasion displaces the homologous parental strand to accommodate the invading transforming ssDNA (hereafter called tDNA). This leads to a three-strand DNA junction, commonly referred to as a displacement loop (D-loop). If the tDNA carries single nucleotide polymorphisms (SNPs) compared to the recipient sequence, its recombination results in an allelic transfer. This phenomenon has led to the hypothesis that, similarity to the role of sexual reproduction in eukaryotes, transformation can help bacteria acquire beneficial mutation or purge deleterious ones (*20–23*). Transformation also leads to sequence acquisition or deletion if a DNA sequence interrupts the region of homology in the tDNA or in the host chromosome, respectively (*24*). The higher efficiency of deletion than acquisition (*25*, *26*) is consistent with the hypothesis that transformation serves a function of curing the genome of deleterious sequence, representing a form of defense against parasitic mobile genetic elements (*21*). Mechanistically, deletion and acquisition involve the formation and resolution of at least two D-loops, one on each region of homology flanking the heterologous sequence. Transformation can produce recombination events, presumably of various lengths, up to several kb-long, involving a dedicated motorization of ssDNA integration at the tips of the transformation D-loop. In monoderm bacteria such as *Streptococcus pneumoniae*, D-loop extension relies on the general recombination helicase RadA (*27*). In contrast, in diderm bacteria such as *Vibrio cholerae* and *Acinetobacter baylyi*, it employs the dedicated transformation helicase ComM which is recruited by DprA onto the D-loop as a ring-shaped hexamer (*28–31*). The helicase activity on parental dsDNA is proposed to promote branch migration around the D-loop, therefore modulating the amount of recombinant tDNA integrated in the chromosome (*26–28*, *30*). Finally, recent advances suggest that chromosome replication contributes to the final resolution of the three-strand intermediates (*28*). However, it remains unclear whether replication alone can resolve recombination D-loops into dsDNA or if this step requires transformation-specific factors.

## YraN and CoiA are two PD-(D/E)XK phosphodiesterases required for natural transformation in distinct species of the bacterial phylum

CoiA, specifically expressed during competence in *S. pneumoniae,* stands out as a candidate possibly involved in resolving the recombination D-loop formed during transformation (*32–35*). CoiA is a cytosolic protein, and detailed genetic indicate that it contributes to events downstream of DNA internalization and protection (*32*, *33*). While displaying an atypical catalytic sequence, CoiA has been classified in the PD-(D/E)XK superfamily of phosphodiesterases (*34*). This raised the hypothesis that CoiA could act as a nuclease at a late step of homologous recombination during natural transformation. To explore the hypothesis that CoiA is a core component of the transformation pathway, we investigated its phylogenetic distribution along with DprA, RadA, and RecA (Fig. 1B and Table S1). RecA, which is usually regarded as ubiquitous across the bacterial phylogenetic tree, was indeed found in 97.5% of the 4,185 queried genomes. The dedicated transformation effector DprA is also widespread (92.6%), only absent in some obligate mutualistic endosymbionts among *Bacteroidota*, mostly *Blattabacterium* and *Buchnera* which are described as also lacking RecA (*36*) (Fig. 1B). Consistent with a role in the transformation-associated recombination, CoiA was inferred to have been acquired within *Bacillota* (formerly *Firmicutes*) which also encode DprA and RecA and was subsequently lost in those lacking RadA.

Thus, considering the hypothesis that CoiA plays a specific and critical role in natural transformation, a functional equivalent is to be expected in the vast majority of CoiA-lacking bacteria. To identify it, we crossed the results of transposon-insertion sequencing (Tn-seq) screens designed to identify all genes required for competence (gene regulation) and natural transformation in the distantly-related diderms *Legionella pneumophila* (*37*) and *Synechococcus elongatus* (*38*) (Fig. 1B). Out of the 13 genes required for transformation and conserved in both species (Table S2), 7 encode the core components of the Type IV retractile filament, 5 encode the highly conserved components of the DNA uptake machinery (ComEC, ComEA, ComFC) and of the homologous recombination pathway (DprA and ComM). The last gene, annotated as *yraN*, encodes a protein of unknown function and belongs to the uncharacterized protein family UPF0102 (PF02021). Like CoiA, YraN harbors the signature structure of PD-(D/E)XK phosphodiesterases, and its catalytic site displays the typical amino acid sequence of this class of enzymes. Its phylogenetic distribution contrasts with that of CoiA: YraN is absent in CoiA-positive *Bacillota* and widely conserved in all other species (Fig. 1B). Indeed, analysis of the patterns of co-occurrence showed that CoiA and YraN are anti-correlated (phyloglm test, effect=-0.73, p=1.5×10^−3^). Reconstruction of the ancestral states of YraN and CoiA in terms of presence-absence indicates that YraN was lost consecutively to the emergence of CoiA in some *Bacillota* (Fig. 1B). Altogether, the results suggest that YraN and CoiA proteins are functional analogues acting similarly in transformation in distinct bacterial species.

To test their contribution to natural transformation, we constructed null mutants of *yraN* in *L. pneumophila* (*Lp*), *A. baylyi* (*Ab*) and *V. cholerae* (*Vc*), and a null mutant of *coiA* in *S. pneumoniae* (*Sp*). Transformation assays using a tDNA with homology to the chromosome and carrying a selectable SNP (allelic transfer event) show that *Lp*, *Ab, Vc,* and *Sp* cells lacking YraN/CoiA can only process a 0.5-5% fraction of transformation events (Fig. 1C to F). Importantly, *yraN* and *coiA* inactivation had no impact on the uptake and establishment of a replicative plasmid via natural transformation (in *Ab*, Fig. S1A, and *Sp*, Fig. S1B). Since plasmid installation does not require homologous recombination in the chromosome, this means that YraN and CoiA are not required in the import steps of transformation and rather have a role in promoting homologous recombination-dependent chromosomal integration of the tDNA. In line with this hypothesis, conjugative transfer of the p45 integrative conjugative element, whose site-specific recombination is expected to be homologous recombination-independent (*39*), is insensitive to inactivation of *yraN* (Fig. S1C). The results are consistent with YraN and CoiA having a role in the specific RecA-directed homologous recombination of tDNA.

Although YraN and CoiA belong to two distinct protein families (PF02021.20 and PF06054, respectively), their display a common fold consisting of four β-sheets sandwiched between two α-helices which form the active site of the large superfamily of PD-(D/E)XK phosphodiesterases (*40*, *41*) (Fig. S2). While YraN shows no other structural domain, CoiA proteins present a small putative C2H2-type zinc finger motif on their N-terminal side and a longer extended region with no structural or functional signature on their C-terminal side. The canonical signature of the phosphodiesterase active site conserved in CoiA and YraN consists of a D residue in β2 and an E residue in β3 (Fig. S2). Expression of YraN mutants of the PD-(D/E)XK active site (YraN^E37A^ and YraN^E50A^ in *Lp*, YraN^E60A^ in *Ab*) did not restore the wild-type phenotype (Fig. 1C and 1D). Immunodetection of a FLAG tag linked at the C-terminus of YraN in *Ab* indicates that transformation defect was not due to YraN destabilization by the point mutations (Fig. S1D). Similarly, the expression of CoiA^D95A^ or CoiA^E105A^ in the *Sp coiA* mutant does not restore the transformation efficiency of a wild-type *Sp* strain (Fig. 1F), while also not altering their transient expression and stability during the competence windows (Fig. S3A). Altogether, these genetic analyses indicate that the phosphodiesterase activity of YraN and CoiA is central to their contribution to the pathway of natural transformation.

## YraN and CoiA are both ssDNA endonucleases, with distinct specificities

PD-(D/E)XK phosphodiesterases define a large and highly diverse superfamily of nucleolytic enzymes with a bipartite catalytic site, divided into several subfamilies characterized by distinct sequence signatures, and by various cleavage specificities modulated either by their target DNA or RNA structure, and/or by their additional domains or protein partners (*40*, *41*). YraN and CoiA belong to two distinct PD-(D/E)XK subfamilies, with YraN harboring the consensus PD-(D/E)XK site and CoiA displaying an atypical site (Fig. S2). To obtain molecular insight into the nucleolytic activities of YraN and CoiA, we analyzed the biochemical properties of soluble recombinant proteins (Fig. S3B and S3D). Gel filtration of purified *L. pneumophila* YraN (YraN*_Lp_*), pneumococcal CoiA and their variants resulted in elution as a single species, in an apparent monomeric form (Fig. S3C and S3E). YraN*_Lp_* displays a nicking activity on dsDNA in a dose-response manner, revealing that it acts as a relaxase on supercoiled circular DNA without further processing (Fig. 2A). As anticipated for this family of phosphodiesterase, this single-strand endonuclease activity requires the E50 residue (Fig. 2A). Importantly, YraN*_Lp_* also processed ssDNA, with highly relaxed site specificity (Fig. 2B). CoiA also acts as a relaxase on a supercoiled plasmid template, without further apparent processing of the DNA (Fig. 2C). However, CoiA displays a higher DNA nicking efficiency than YraN*_Lp_* when assayed with the same gradual range of protein concentration. In stark contrast to YraN*_Lp_*, CoiA barely processed the linear ssDNA template regardless of protein concentration (cleavage of 2% of total substrate, Fig. 2D). We hypothesize that CoiA activity relies on its interaction with dsDNA to nick one strand of the supercoiled plasmid template. This would mean that CoiA acts on a specific sequence or structure, which distinguishes if from YraN.

**Fig. 2:**
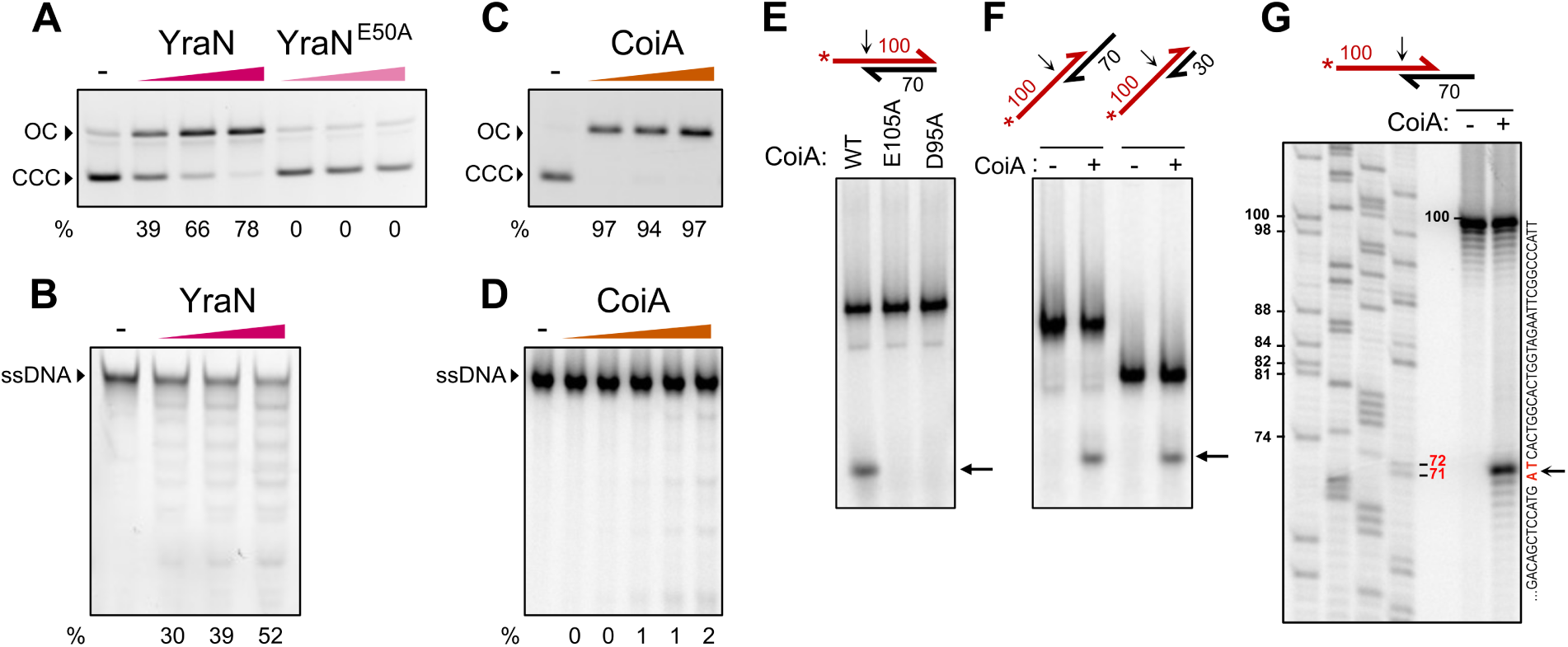
YraN and CoiA are ssDNA nucleases, with CoiA specific to 5’-3’ ssDNA-dsDNA junctions. (A) Agarose gel electrophoresis of plasmid DNA incubated with increasing amount (wedges) of YraN*_Lp_* (∼200; 400; and 800 nM) and E50A mutant (∼300; 600; and 1200 nM). Incubation with YraN*_Lp_*, but not with the mutant E50A, produces a shift of the supercoiled fraction (Covalently Closed Circle, CCC) to the relaxed form (Open Circle, OC), indicative of ssDNA phosphodiesterase activity. (B) Acrylamide gel electrophoresis of fluorescent (Cy3) 100-mer ssDNA incubated with increasing amount of YraN*_Lp_* (∼200; 400; and 800 nM), after protein removal. The ladder formed below the main ssDNA band reveals endonuclease activity. (C) Agarose gel electrophoresis of plasmid DNA incubated with increasing amount (∼150; 300; and 600 nM) of CoiA. (D) Nuclease assays performed with increasing amount of CoiA (∼8; 30; 80; 300; and 800 nM) on a radiolabelled (^33^P) 100-mer ssDNA fragment, migrated on PAGE after protein removal. Values of the relative amounts of cleaved products (in % of total DNA) in each assay are reported below the gel. (E)(F) Nuclease assays visualized on PAGE after protein removal: (E) Activity of CoiA (WT) and its derivatives (E105A, D95A) and (F) Activity on various substrates with (+) or without (-) CoiA (WT). Template DNA structures are schematized on top of each panel and detailed in Figure S7. The numbers indicate the size in nt of the oligonucleotides. Half arrow extremities symbolize 3’-ends of DNA strands. Black arrows point to CoiA cleavage sites. The 5’-radiolabelled (*) oligonucleotide is depicted in red. The black arrow next to the gels show the band resulting from substrate cleavage by CoiA. (G) Mapping of CoiA-dependent (∼300 nM) DNA cleavage site (vertical right) on denaturing sequencing gel using unrelated DNA sequencing reactions as nucleotides ladder (left).

## CoiA is structure-specific endonuclease

To characterize CoiA endonucleolytic specificity, we analyzed its interaction and cleavage activities on various linear DNA templates comprising dsDNA and/or ssDNA segments of different lengths and polarity. Electromobility shift assays (EMSA) performed with a 100 nucleotide-long radiolabeled linear DNA substrate showed that CoiA interaction is more stable with ssDNA than with dsDNA, indicating that the lack of cleavage of the ssDNA does not correlate with a lack of interaction (Fig. S4A). CoiA binding on ssDNA generates a ladder-like shift pattern which depends on the length of the template, as observed with the two dsDNA substrates extended by a 3’ or a 5’ ssDNA tail of different sizes (Fig. S4A). Interestingly, an additional and faster migrating DNA species was observed upon interaction of CoiA with the 5’-tailed DNA substrate (Fig. S4A) that we could better analyzed after protein denaturation (Fig. 2E). This additional labeled DNA species was absent with two purified point CoiA mutants of consensual positions of PD-(D/E)XK motif, *i.e.* the E105A and D95A mutants (Fig. 2E). In all, these experiments indicate that CoiA catalyzes a single and precise endonucleolytic cleavage in this 5’-tailed dsDNA substrate, at a position that matches with the ss/dsDNA junction.

We further investigated this endonucleolytic reaction with other linear DNA templates. We used a 30 nts-long dsDNA substrate, extended by a 5’ tail of 70 nts on one side and 40 nts on the other, thereby forming two 5’ ss-dsDNA junctions. Intriguingly, CoiA cleaved this dual 5’-tailed DNA template only at one ss-dsDNA junction, but not at the other (Fig. 2F). This cleavage was also obtained using the same dsDNA substrate without the 40 nts-long 5’ ssDNA tail (Fig. 2F). CoiA cut the radiolabelled DNA strand within the dsDNA moiety of the template at the −1 position of the ss-dsDNA junction, between an A and T (Fig. 2G). These results suggest that CoiA exhibits a sequence specificity to cleave such junctions. This indicates a preferred cutting site between an A and a T. To inquire in more detail this cleavage specificity, we generated nine 5’-tailed DNA templates differing on the 4 nts overlapping the ss-dsDNA junction, *i.e.* 1 nt in the ssDNA part and 3 nts in the dsDNA part and we also generated one DNA hybrid structure differing in the ssDNA tail orientation. It appears that CoiA cleaves the tailed strand of a ss-dsNA junction preferentially made of 5’ GATC 3’ sequence, between the A and T bases in the duplex DNA (Fig. 4B and S4B). Of note, CoiA did not cut a 3’-tailed dsDNA made of this preferred DNA sequence at the ss-dsDNA junction (Fig. S4B, structure 10).

Altogether, these biochemical analysis of YraN and CoiA demonstrate that both are ssDNA endonucleases, relying on their PD-(D/E)XK motif to cleave DNA. YraN appears to cut ssDNA with little sequence and structure specificities. In contrast, CoiA is revealed to be a Structure-Specific Endonuclease (SSE). Indeed, CoiA acts on a ss-dsDNA junction of a 5’-tailed dsDNA, at the −1 position within the dsDNA portion.

## CoiA is a recombination D-loop resolvase

SSE enzymes are involved in many different processes of DNA dynamics in all kingdoms of life (*42*, *43*). In particular, they include nucleases that process branched DNA intermediates of homologous recombination, which are commonly referred to as resolvases (*43*).

We hypothesized that CoiA could act as a resolvase on the three-stranded DNA intermediate generated in the particular ssDNA recombination pathway of natural transformation. To test this, we first assessed CoiA nucleolytic activity on several branched DNA structures assembled with partially complementary synthetic oligonucleotides, including its preferred GATC ss-dsDNA cleavage junction characterized on linear DNA. Four distinct structures were tested: a flayed DNA duplex, a 5’ flap DNA, a DNA bubble, and a D-loop-like structure. CoiA was able to cleave all these substrates (Fig. 3A), and precise mapping of the two latter DNA substrates showed that CoiA cleavage happened exclusively at the 5’ss-dsDNA junction (as represented in Fig. 3B). CoiA processed these four DNA structures with nearly identical efficiency, with a small preference for the 5’ flap and the D-loop-like templates (Fig. 3A).

**Fig. 3:**
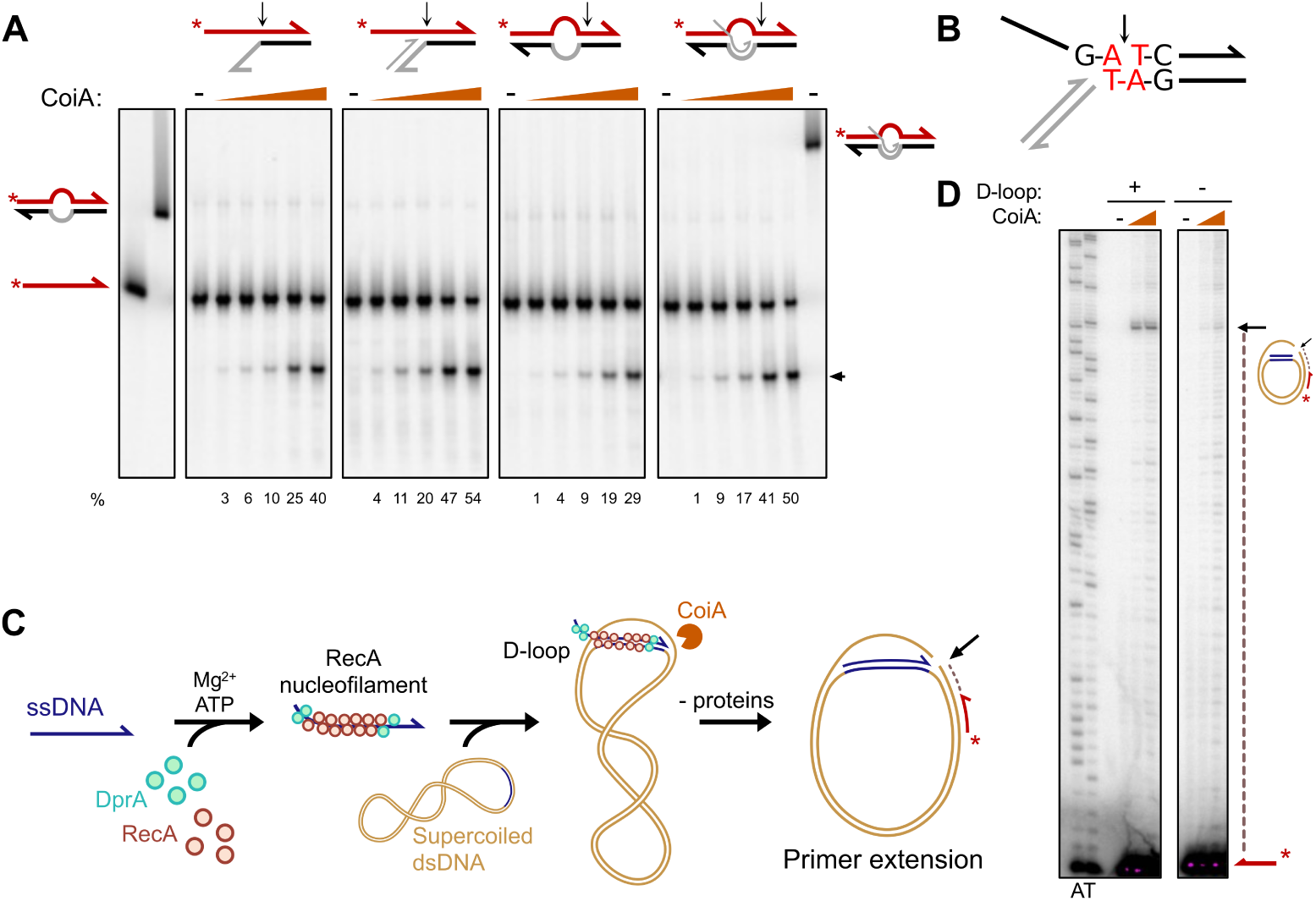
CoiA is a resolvase that recognizes and cleaves D-Loop junctions. (A) Nuclease assays of increasing amounts of (∼8; 30; 80; 300; and 800 nM) on various branched DNA structures, migrated on PAGE after heat denaturation of DNA strands and protein removal. Template DNA structures are schematized on top of each panel and detailed in Fig. S7. Half arrow extremities symbolize 3’-ends of DNA strands. The 5’-radiolabelled (*) oligonucleotide (100 nt) is depicted in red. Black arrows point to CoiA cleavage sites. Values of the relative amounts of cleaved 5’-radiolabelled products (% of total DNA) are reported below the gels. A dsDNA bubble, a ssDNA probe, and a D-loop structure without heat denaturation were used as migration markers to check the quality of structures and the denaturation of the DNA strands and are depicted on the sides of the gel. (B) Representation of the CoiA DNA cleavage site on branched DNA. (C) Schematics of the experimental procedure of the D-loop nucleolysis assay. CoiA nuclease activity is tested on a D-Loop reconstituted on a supercoiled dsDNA plasmid (gold) with homologous ssDNA oligonucleotides (homology in blue), pre-incubated with RecA, DprA, ATP, and Mg^2+^, to form a RecA nucleofilament. (D) D-loop nucleolysis assay. Mapping of the cleavage at the D-Loop extremity (black arrow) after protein removal, using a 5’-radiolabelled (*) primer (red) extension experiment (dotted line) and migration on urea sequencing PAGE. Experiments are performed without (-) or with increasing amount of CoiA (125; 375 nM). As a control, this experiment was also performed with a non-homologous ssDNA (D-loop -). Chain-Termination DNA sequencing reaction with ddA (A) and ddT (T) using the same primer on the plasmid were loaded in the same gel (left). See Fig. S5 for control experiments.

CoiA ability to cleave an artificial D-loop-like structure strongly supports the proposal that CoiA can act as a homologous recombination D-loop resolvase during transformation. However, the unpaired ssDNA strand is not identical to the paired DNA strand in this artificial three-strand DNA structure. This markedly differs from a native D-loop assembled by RecA in which the invading strand is identical to the extruded strand. In addition, RecA and other homologous recombination effectors might also modulate the accessibility and/or cleavage activity of CoiA to the D-loop intermediate. Thus, to investigate CoiA nucleolytic activity on a *bona fide* homologous recombination D-loop, we set up an *in vitro* assay relying on the concerted actions of purified pneumococcal RecA and DprA, the two dedicated homologous recombination mediators of transformation (*18*, *19*) (Fig. 3C). First, we optimized the experimental conditions to efficiently convert supercoiled DNA plasmid molecules into D-loop structures via DprA-mediated and RecA-directed ssDNA strand exchange reaction (Fig. S5A (*44*). This resulted in appropriate *in vitro* conditions to analyze the nucleolytic action of CoiA on a D-loop structure as assembled during transformation. We used a 100 nt-long homologous ssDNA oligonucleotide that generate the preferred 5’ ss-dsDNA CoiA cleavage site when fully paired at its 3’ end with the complementary sequence in the supercoiled plasmid (Fig. 3C). CoiA added to this preformed homologous recombination D-loop cleaved the DNA: these cleavage sites were mapped by run-off as depicted in Fig. 3D. CoiA preferentially and specifically cleaved the displaced DNA strand of the D-loop at the exact position expected for the homologous pairing of the ssDNA oligonucleotide at its 3’ end in the plasmid (Fig. 3D). Other cleavages of weaker intensity were detected upstream and downstream the main cleavage site at the boundary of the D-loop. A control experiment performed with a heterologous oligonucleotide demonstrated that this main cutting site in the plasmid paired with the homologous oligonucleotide resulted from CoiA cleavage at the 3’ boundary of the D-loop (*i.e.*, corresponding to the 3’ end of the paired oligonucleotide) (Fig. 3D and S5B). In addition, the same weaker DNA cleavages were detected in this control experiment, which probably resulted from CoiA relaxase activity on the supercoiled plasmid. Altogether, these results show that CoiA is a homologous recombination D-loop resolvase specifically cleaving the displaced strand at the three-strand DNA junction located at the 3’ side of the D-loop.

## CoiA and YraN act synergistically with the RadA and ComM helicases, respectively

Another key pneumococcal transformation effector involved in the maturation of transformation D-loop is RadA, a widely conserved bacterial helicase promoting DNA branch migration of the three-strand junctions of the D-loop to extend integration of the invading ssDNA (*27*). This led us to analyze the epistatic relationship between CoiA and RadA in transformation.

First, transformation assays with a chromosomal tDNA selective for an allelic transfer event showed similar 30- to 50-fold reduction in transformation efficiency for *coiA*, *radA*, and *coiA radA* null mutants (Fig. 4A). This suggests the involvement of CoiA in RadA-mediated D-loop processing. We previously designed an experiment to test RadA helicase activity in transformation (*i.e.*, extending ssDNA incorporation in D-loops) using PCR fragments of similar size (∼4 kb) carrying a selectable SNP located either centrally (PCRc) or close to one end (PCRe) (*27*). Due to the random cleavage of these fragments at the initial stage of transformation, and because they are internalized by their 3′ end, the selectable SNP is equally distributed between the middle and the 3′ end of the transforming ssDNA from PCRc. However, for PCRe, the SNP is located almost exclusively near the 5′ end, increasing the requirement for D-loop extension for its acquisition. We found that, similarly to RadA, CoiA absence impairs transformation to a greater extent for PCRe compared to PCRc (122-fold in the *radA* null mutant and 25-fold in the *coiA* null mutant, Fig. 4B). Notably, RadA appears to be more important than CoiA for PCRe transformation efficiency. Since the transformation defects of the *radA* and *coiA* mutants are not cumulative, this suggests that CoiA and RadA act together synergistically, defining a helicase/nuclease system processing the D-loop formed during transformation.

**Fig. 4.**
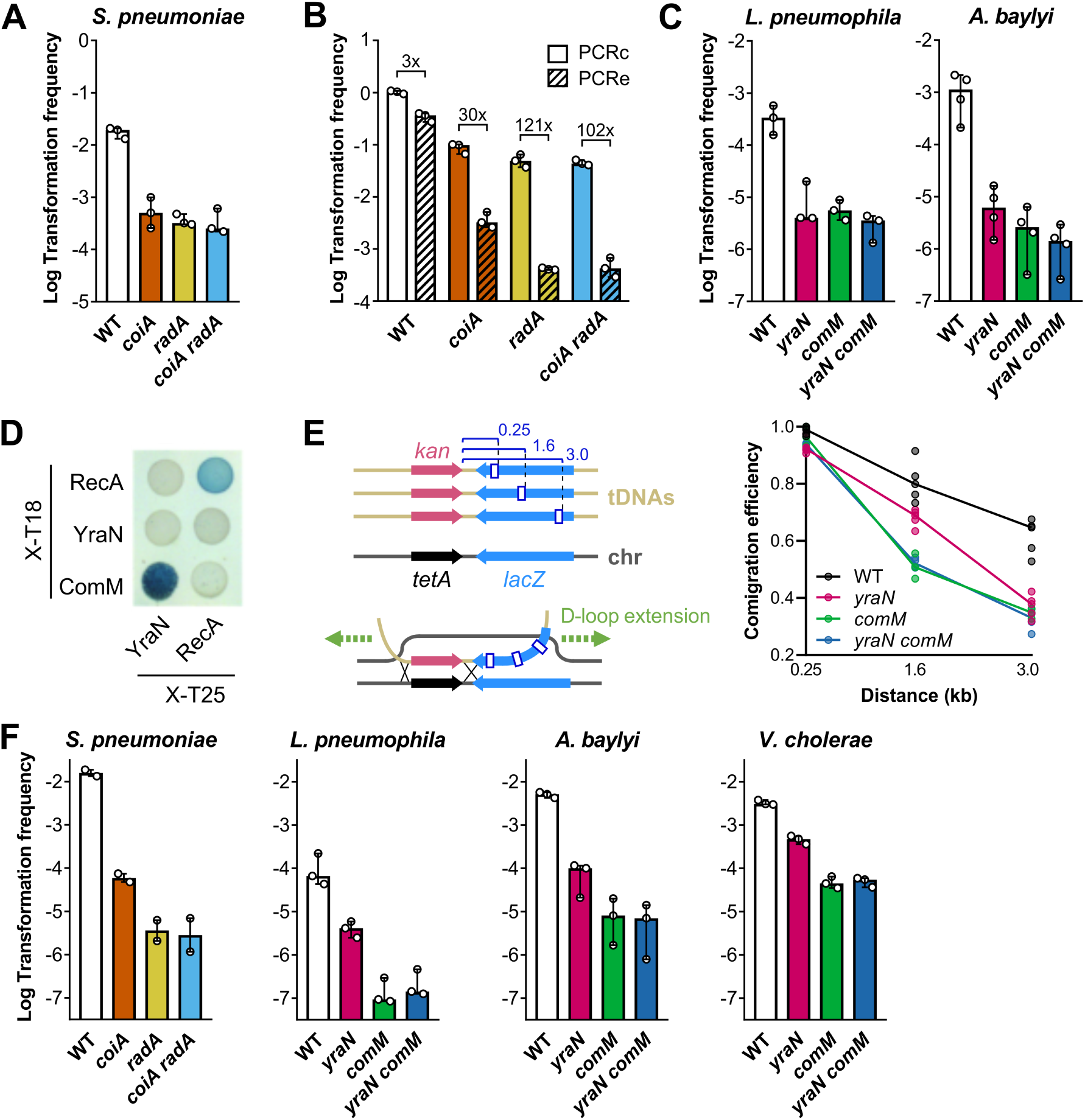
YraN and CoiA are involved in ComM- and RadA-mediated D-loop extension. (A) Transformation frequency of *Sp* parental (WT) and mutant strains transformed with the parental gDNA carrying a Rif^R^-conferring point mutation as tDNA. Bars represent the median with range of three independent experiments. (B) Transformation frequency of *Sp* strains with 4 kb-long PCR products amplified from strain R304 chromosomal DNA and carrying the Rif^R^-conferring point mutation either centered (PCRc, plain bars) or off-center (PCRe, hatched bars). Bars represent the median with range of three independent experiments. (C) Efficiency of transformation-mediated allelic transfer of *Ab* and *Lp* parent and mutant strains. Cells were exposed to a tDNA PCR product consisting of either a Rif^R^-conferring *rpoB* allele (*Ab*) or a Strep^R^-conferring *rpsL* allele (*Lp*). Bars represent the median with range from three and four independent experiments. (D) BACTH to detect interaction between the *Lp* YraN, RecA, and ComM proteins. (E) Left: Principle of the *in vivo* branch migration assay based on marker comigration. An *A. nosocomialis* strain carrying a tetracycline resistance gene followed by the *lacZ* gene (chr, chromosome, grey strand) is exposed to a tDNA (gold strand) carrying a kanamycin resistance gene (*kan*) instead of the tetracycline cassette, along with a 7-bp deletion in the *lacZ* gene at 0.25, 1.6, or 3 kb from the end of the kanamycin gene. Right: Comigration marker experiment. Comigration efficiency (Kan^R^ lacZ^-^ CFU *vs* all Kan^R^ CFU) are determined as a function of the distance separating the markers in WT and mutants of *A. nosocomialis*. Lines correspond to the average of five experiments (individual dots).(F) Efficiency of transformation of an insertion event in *Sp*, *Lp*, *Ab,* and *Vc*. The tDNA respectively corresponds to parental gDNA carrying the *comX* gene interrupted by a tetracycline resistance gene, a PCR product of a chromosomal region (*lpp2553*) interrupted by an apramycin resistance gene, and a PCR product carrying a kanamycin resistance gene corresponding to the *kan-lacZ* cassette used in (E), and a PCR product of the VC1807 gene interrupted by an erythromycin resistance gene. Both PCR products are homologous to the recipient DNA over 2 kb on both sides of the cassette. Bars represent the median with range of two or three independent experiments.

We next asked whether the same synergy applies to YraN and ComM, the helicase dedicated to the extension of the transformation D-loop in diderms. A functional association is supported by the phylogenetic distribution of the *comM* and *yraN* genes (Fig. 1B), with the presence of *comM* positively associated with the presence of *yraN* (phyloglm test, effect=4.16, p=8.25×10^−4^). The genes are co-localized in the genome of 34% of the bacterial genomes that have both *comM* and *yraN*, which is often a sign of functional interaction. We measured the transformation frequency for an allelic transfer event in *yraN* and *comM* mutants of *Lp* and *Ab*. This transformation assay showed reduction in transformation frequencies for both mutants in both species (Fig. 4C). Furthermore, these mutations are not cumulative as the *yraN comM* double mutants exhibit a similar transformation defect, clearly suggesting that YraN and ComM cooperate in the transformation pathway.

Two large-scale co-evolution analyses previously predicted the direct interaction of these proteins (*45*, *46*). We thus tested these interactions using the bacterial two-hybrid assay based on the functional reconstitution of the *Bordetella pertusis* adenylate cyclase from its T18 and T25 domains fused to the bait and prey proteins (*47*). Validating the assay, RecA is found to self-interact (Fig. 4D) (*48*). No interaction between RecA and ComM has been detected, but we did observe the predicted interaction between YraN and ComM (Fig. 4D). To further genetically test the hypothesis that YraN might assist ComM in D-loop extension, we determined the efficiency of integration of genetic markers separated by increasing distances (comigration marker experiment). An *Acinetobacter nosocomialis* strain carrying an integrated tetracycline resistance gene followed by the *lacZ* gene was exposed to transforming DNA encompassing the tetracycline resistance, but replaced by a kanamycin resistance gene, and with a 7-bp deletion in the *lacZ* gene at an increasing distance from the kanamycin resistance marker as illustrated in Fig. 4E. Selecting transformation events using the kanamycin marker, we determined the comigration efficiency of the unselected 7-bp deletion (Kan^R^ *lacZ*^-^ CFU vs all Kan^R^ CFU) as a function of the distance separating these markers. In the wild-type background, the comigration efficiency decreases with the distance separating the markers, from nearly 1 at 0.25 kb distance down to 0.7 at 3 kb (Fig. 4E). While comigration at 0.25 kb distance is unchanged in a ComM-deficient strain, it rapidly decreases with increasing distance, down to 0.4 at 3 kb. This confirms previous results obtained in *V. cholerae* and *A. baylyi* (*30*). The *yraN* mutant is also severely affected for comigration at 3 kb, similar to the *comM* mutant, suggesting that *yraN* is required for the ComM-dependent D-loop extension. Yet, at 1.6 kb, the *yraN* mutant is less deficient than the *comM* mutant, suggesting that YraN facilitates ComM-dependent D-loop extension over long distances. Consistent with results from transformation assays, the deficiencies in comigration of the *comM yraN* double mutant are not additive. Altogether, the results indicate that ComM and YraN function as a helicase/nuclease system dedicated to the maturation of the transformation D-loop.

For both RadA/CoiA and ComM/YraN helicase/nuclease systems, the nuclease mutant is less severely impaired than the helicase mutant in assays testing for D-loop extension. This suggests that the nuclease is mainly required to extend the D-loop while its initiation and stabilization rely primarily on the helicase. To test this hypothesis, we measured the transformation efficiency of an event resulting in an insertion, using a tDNA carrying an antibiotic cassette (Fig. 4F). Such event involves the homologous recombination of both regions flanking the heterologous sequence, hence requiring the formation of at least two D-loops. Unlike the branch migration assay, the success of this incorporation may not strongly depend on the extension of these D-loops. As expected, mutants of the ComM helicase show a strong reduction in transformation frequencies (≥ 2-log reduction compared to the wild-type, Fig. 4F), consistent with previous observations (*26*, *30*). Similarly, the RadA mutant is severely defective (∼4000-fold reduction compared to the wild-type). In all tested species, mutants of the nuclease are also deficient in transformation, yet less so than the mutants of the helicase. The transformation deficiencies of the helicase mutants are dominant over, but not additive to, those of the nuclease mutants. These results are consistent with a model in which the YraN and CoiA nucleases process the D-loop in a way that facilitates the D-loop extension by their cognate helicase.

## The RadA/CoiA and ComM/YraN helicase/nuclease systems differ in promoting extended recombination events

To further characterize the role of ComM/YraN and RadA/CoiA helicase/nuclease pairs in the integration of transforming DNA, we sought to obtain a high-resolution map of recombination events from natural transformation. To this aim, we exploited the fact that a sequence with multiple mismatches to the native homolog (homeologous sequence) could be recombined by the transformation pathway. While a single selectable SNP can provide a quantitative readout of allelic transfer efficiency, we sought to track the fate of several unselected SNPs in order to generate a high-resolution view of the successful recombination events (Fig. 5A). However, the recipient DNA mismatch repair system (MMR) could recognize and correct mismatches generated during heteroduplex formation in the newly transformed cells, therefore erasing the informative recombination markers. To prevent this, we used a *mutS* null mutant of *Ab* (MMR^-^, annotated as wild-type for this part of the manuscript) and a *hexA* null mutant of *Sp*. As tDNA sequence, we selected the RNA polymerase-encoding loci: in closely related species, gene synteny is maintained in this region which contains essential genes involved in transcription and translation. We transformed wild-type and mutant strains of *Ab* with a 6.4 kb-long PCR fragment encompassing the *rpoB* gene from a rifampicin-resistant isolate of *A. nosocomialis* M2. This homeologous fragment differs from the *Ab* recipient sequence by 841 SNPs with an average of 1 SNP every 8 bp (Fig. 5 and S6). In *Sp*, we used a 6.4 kb Rif^R^-conferring sequence from *Streptococcus mitis* differing from the *Sp* sequence by 251 SNPs, with an average of 1 SNP every 25 bp.

**Fig. 5.**
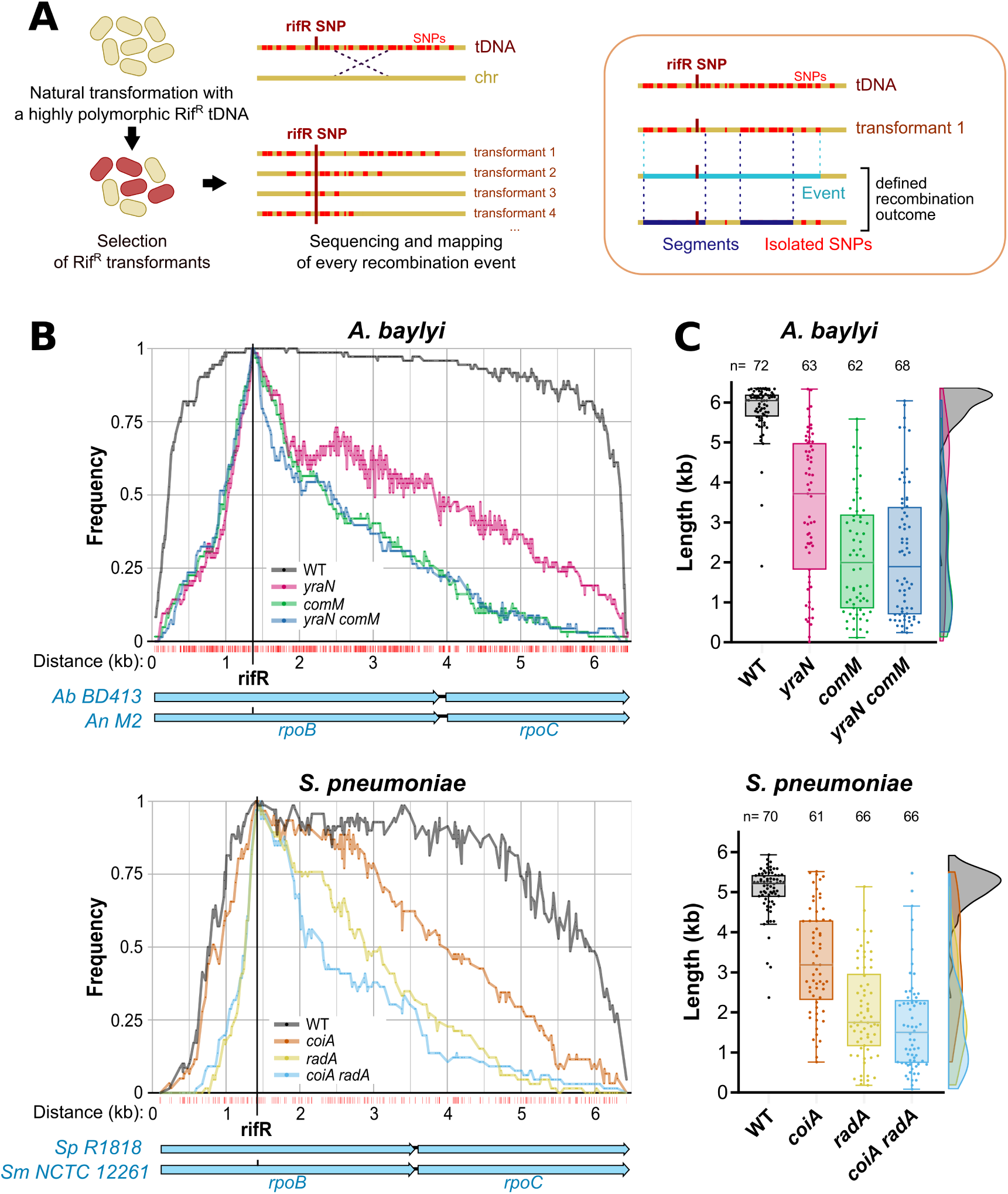
YraN/CoiA promote integration of transforming DNA in association with the ComM/RadA helicases. (A) Left: Schematic representation of the recombination events mapping assay. Parental and mutant strains of *Ab* (*mutS* null) and *Sp* were naturally transformed with a 6.4 kb-long homeologous PCR fragment (gold with red mismatches) carrying an off-centered rifampicin resistance-conferring SNP (rifR, red vertical bar). PCR fragments were amplified from *A. nosocomialis* M2 (carrying 841 SNPs relative to the *Ab* recipient region), and from *S. mitis* (carrying 251 SNPs relative to the *Sp* recipient region). Site-specific sequencing of 62 to 72 transformants of each strain is used to determine the acquired SNPs, defining recombination events. Right: Illustration of metrics terms used in the manuscript. Individual SNPs (red) is the raw data obtained by sequencing. “Event” (light blue) refers to the distance between the two distal recombinated SNPs. “Segment” (dark blue) refers to a continuous uninterrupted stretch of at least two recombinated SNPs (perfectly homologous to the donor sequence). (B) Frequencies of acquisition of each SNP over the 6.4 kb sequence during the recombination events mapping assay in the parental and mutant strains of *Ab* (top) and *Sp* (bottom). The frequency corresponds to the number of samples where each position was acquired, normalized with the total number of samples for the given strain. The selected off-centered a Rif^R^-conferring SNP (therefore present in 100% of the transformants) is represented by a vertical red line. The data was acquired over three independent experiments. See Fig. S6 for detail in every transformant. (C) Quantitative analysis of the length of recombination events observed in panel (B) for *Ab* (top) and *Sp* (bottom). Recombination event length corresponds to the distance separating the two distal SNPs acquired in each transformants (as illustrated in (A)).

In *Ab*, inactivating *yraN*, *comM*, or both, resulted in around 2-log transformation defect of the Rif^R^ homeologous DNA compared to the wild-type strain (Fig. S6A). Similarly, a 2-log reduction of transformation efficiency is observed in the *coiA*, *radA*, and *coiA radA* mutants of *Sp* when transformed with the homeologous Rif^R^ DNA (Fig. S6A). This strong reduction in transformation frequency indicates that the recombination events resulting in selectable Rif^R^ transformants in the mutants represent only a fraction of the events that occurred in a wild-type strain. This can represent an inherent bias in comparing the characteristics of the recombination events, underestimating possible differences between wild-type and mutant strains. We sequenced the targeted region and mapped the acquired SNPs resulting from the recombination events in 62 to 72 transformants of each genotype (wild-type, *yraN*, *comM*, and *yraN comM* in *Ab*; wild-type, *coiA*, *radA*, and *coiA radA* in *Sp*). While all transformants acquired the selected *rifR* SNP, the acquisition frequency of unselected SNPs decreases with the distance separating them from the selected SNP (Fig. 5B). In both *Ab* and *Sp*, the decline in SNP conversion rate around the *rifR* SNP is more pronounced in the helicase and nuclease mutants compared to their respective wild-type strains (Fig. 5B). This observation corroborates the results obtained in the comigration genetic assay (Fig. 4E).

We measured the length of the recombination events, defined as the sequence stretching from the two outermost acquired SNPs (as illustrated in Fig. 5A). Median recombination event length is 5.2 kb in wild-type *Sp* transformants and 6.0 kb in *Ab* transformants (Fig. 5C). Nuclease and helicase mutants show significantly shorter events compared to their cognate wild-type strains (Games-Howell test, p-value < 0.05) with a median length of 3.2 kb and 1.8 kb for *coiA* and *radA* mutants, respectively, and 3.7 kb and 2.0 kb for *yraN* and *comM* mutants, respectively. Since the mutants produce shorter recombination events, their transformation defect is likely due to fewer cells acquiring the selective SNP despite the same number of tDNA invasions as in wild-type strains. In other words, a reduced capacity for D-loop extension decreases the probability of obtaining the selective SNP within the population that recombined DNA. Interestingly, the *radA* and *coiA radA* mutants show identical (median of 1.8 and 1.5 kb-long events respectively) and stronger defects than the *coiA* mutant (Games-Howell test, p-value < 0.05). In accordance with the comigration assay, the same observation is true in *Ab*: event length of *comM* and *yraN comM* mutants are identical (median of 2.0 and 1.9 kb-long events respectively) and shorter than those of the *yraN* mutant (Games-Howell test, p-value < 0.05). These results indicate that the nuclease cannot contribute to DNA integration in the absence of the helicase, therefore suggesting that YraN/CoiA nucleases act as a facilitator of ComM/RadA helicase activity (Fig. 5C). Such similar phenotype in the two tested species is remarkable, highlighting a conserved mechanism in distantly-related species and based on distinct molecular components.

We nonetheless noted differences in the two species regarding how the marker SNPs are acquired or not, generating more or less fragmentation in recombination events. Since this may be due to specificities of the helicase/nuclease systems, we further analyzed the recombination events. To do so, we defined as recombination “segments” the continuous regions of more than one acquired SNP which are interrupted by one or several non-acquired SNPs (Fig. 5A). Non-consecutive SNPs are thus considered isolated recombined SNP. The recombination events of wild-type *Sp* were highly fragmented with 1 to 45 segments and SNPs per event, while in wild-type Ab, the recombination events showed fewer interruptions, with only 1 to 15 segments and SNPs per event (Fig. S6B and C). Since we used MMR-deficient strains, this fragmentation is not due to the reversion of the acquired SNP into the original sequence by mismatch repair. An alternative explanation is that one or more tDNA molecules (possibly fragmented during uptake) can create multiple D-loops, each being stabilized and processed by the helicase/nuclease system. High rates of D-loops processing, even if not extended, would result in apparent branch migration, as observed in Fig. 4. Stochasticity in tDNA invasions and D-loop stabilization/processing can thus produce randomly fragmented events. The segments in *Sp* are not significantly shorter in mutants lacking CoiA/RadA, consistent with the observed recombination events being the product of the helicase/nuclease system stabilizing/processing multiple D-loops with little D-loop extension (Fig. S6D). In contrast, in *Ab*, all mutants display significantly shorter segments than the wild-type strain (Dunn’s test, p-value < 0.05, median of 0.8 kb vs. 5.4 kb, Fig. S6D), suggesting that efficient D-loop extension by the ComM/YraN system allows more frequent merging of D-loops, producing continuous recombination events

In conclusion, although the mechanism of D-loop recognition/cleavage by the nuclease and the nature/processivity of the helicase might differ, these helicase/nuclease systems promote natural transformation by increasing the rate and the extent of incorporated tDNA.

## Discussion

Nearly a century after the description of natural transformation by Frederick Griffith (*49*), several key steps of the process remain unresolved, in particular how ssDNA is stably integrated in the chromosome. The mechanism of homologous recombination relying on a linear ssDNA is unique to the transformation process and might differ from other recombination pathways, that all involve a dsDNA moiety in the two exchanged DNA templates. We propose a model for the integration of tDNA involving two distinct but functionally equivalent helicase/nuclease (Fig. 6).

**Fig. 6.**
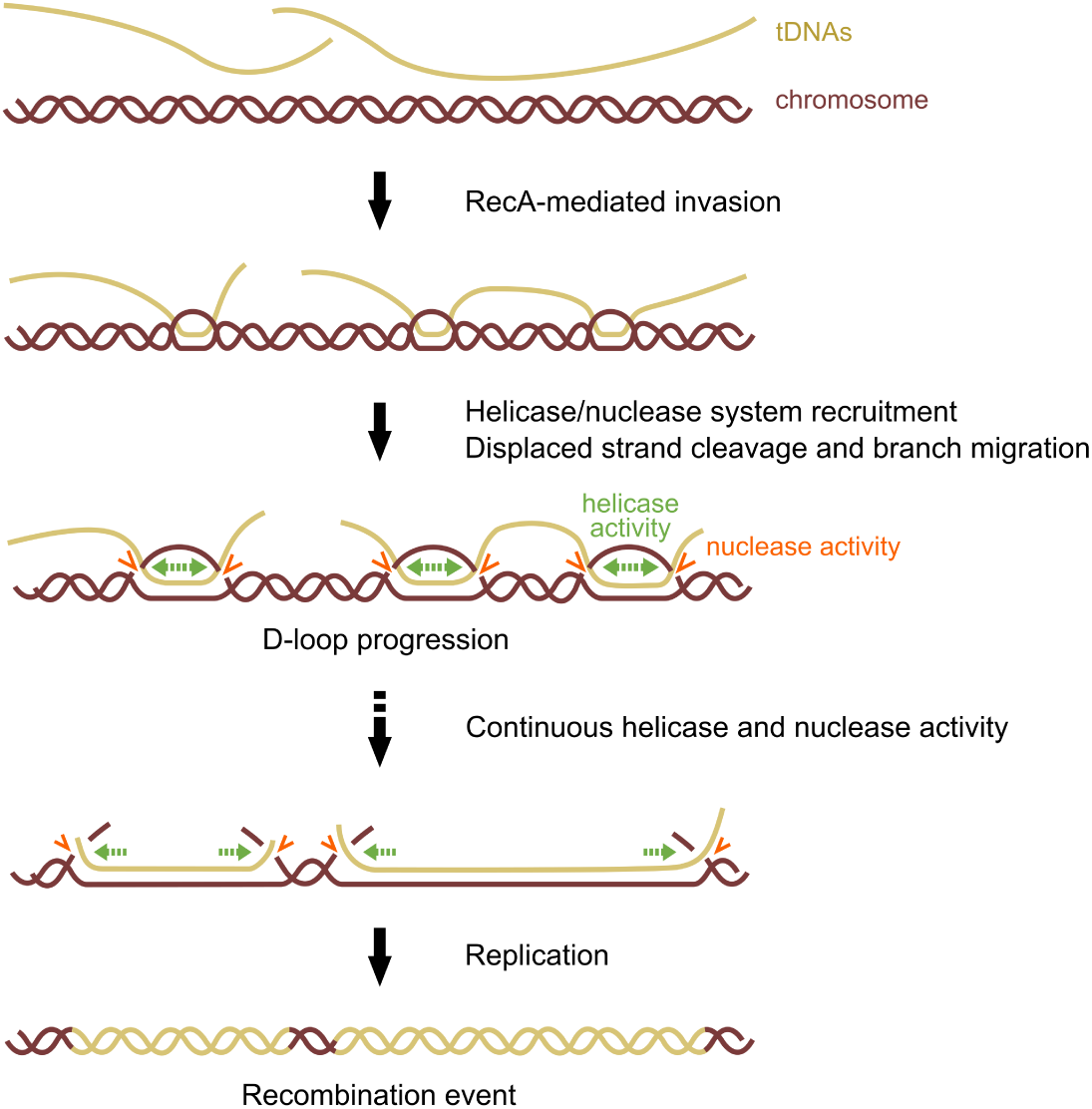
Model for nuclease contribution to recombination during natural transformation. Recombination D-loop formation. The parental strand is displaced by the invasion of the chromosome (dark red) by the tDNA (gold) mediated by RecA (not shown). This step is reversible by spontaneous RecA dissociation. Once the helicase/nuclease system is recruited, helicase activity of ComM or RadA allows D-loop extension (green arrows), while YraN or CoiA nuclease nicks the displaced strand (orange arrows). We suggest that D-loop progression is irreversible after nicking of the displaced strand, which would prevent helicase backtracking and subsequent re-annealing of the two parental strands. Such reversion might happen in nuclease mutants. During branch migration, the D-loops can merge to form a continous recombination event. D-loop resolution occurs before or during replication through an unknown process.

After its entrance into the cell, transforming ssDNA is bound by DprA and RecA, shielding it from nuclease activity (*18*). Following its nucleofilamentation mediated by DprA, RecA displaces one chromosomal strand to form a heteroduplex between the invading tDNA and the complementary chromosomal strand. This creates a D-loop structure which is next processed, depending on the species, either by the generic DnaB-type RadA helicase, recruited via its interaction with RecA, or by the transformation-specific MCM-type ComM helicase via its interaction with DprA, thereby providing specificity toward the transformation D-loop. The helicase then unwinds the chromosomal DNA and promotes the extension of the D-loop (*27*, *28*, *31*). Our results show that YraN or CoiA nuclease also act on and maturate the D-loop, synergically with their cognate helicase, by ssDNA endonucleolytic cleavage to favor the integration of the invading strand at the expense of the displaced parental strand (Fig. 6). We hypothesize that YraN is recruited and/or activated through direct interaction with ComM (Fig. 4D). This process would be similar to that of CRISPR and argonaute systems, with the helicase directing the nuclease towards cleavage of the displaced strand (*50*). This would ensure that YraN, which shows activity against any ssDNA, is targeted at the transformation D-loop where the displaced strand might be exposed because neither involved in a duplex nor protected by DprA like the tDNA strand. In contrast, CoiA is an SSE recognizing and specifically cleaving branched DNA structures at a ss-dsDNA junction. CoiA can thus specifically cleave the displaced strand of the transformation D-loop (Fig. 3D), a feature that may avoid spurious cleavage of ssDNA outside of transformation. For both YraN and CoiA, cleavage of the displaced strand might not be (or loosely) sequence-specific (as suggested by Fig. 2B).

How could cleavage of the displaced strand promote the efficient integration of transforming DNA? Cleavage of the displaced strand by YraN/CoiA could prevent helicase backtracking and reversion of the D-loop. Only about 25% of invasion events followed by the recruitment of ComM successfully produced recombinants (*28*), suggesting that strand invasion and initial D-loop extension are reversible events. The nuclease would thus have a role in D-loop stabilization, resulting in a higher success rate of recombination events. In addition, the nucleases can act as relaxases to ease D-loop extension during DNA unwinding. With the need for cleavage arising in the context of helicase-driven D-loop progression, this implies that D-loops are actively extended. Yet, it is unclear how recombination events are formed and how much they depend on the extension of D-loops. The analysis of recombination events, based on the detection of SNPs acquisition, leads to the hypothesis that extended recombination events emerge from several invasions of one or more tDNA molecules. Indeed, transformation-acquired mosaicism was previously reported in several species, where several recombined segments are thought to arise from a single tDNA molecule or through simultaneous transformation events (*51–53*). We propose that the difference in the nature of the recombination events, (continuous in *Ab*, more fragmented in *Sp*) arises from the extent to which the multiple D-loops formed by tDNA invasions are extended by the helicase/nuclease systems. The limited D-loop extension paired with high fragmentation in wild-type *Sp* hints that the main role of the RadA/CoiA system is D-loop stabilization with limited D-loop extension. It is possible that extensions are prevented by the mismatches. In *Ab*, longer and more continuous recombination events suggest that YraN/ComM is important for D-loop extension and can bypass mismatches. This also implies a higher processivity or stability of ComM on DNA compared to RadA, although such properties remain to be demonstrated. The helicase/nuclease pairs can thus produce extended recombination through both D-loop stabilization and extension, where the nuclease role is to assist its cognate helicase. Interestingly, in the absence of the helicase, many short recombination events are still produced (Fig. 5C), albeit at about 100-times less (Fig. S6A). This indicates an unknown and less processive alternate D-loop processing pathway, possibly mediated by RecA and accessory proteins acting in DNA repair pathways of genome maintenance, to which the nuclease cannot assist. Nonetheless, when both helicase and nuclease are present, their action appears to be deeply coordinated, and the molecular mechanisms behind their synergistic action will warrant biochemical and structural biology investigations.

Why would bacteria have evolved helicase/nuclease systems to foster recombination by transformation? We found that the transformation helicase/nuclease pairs act to reduce the stretches of unacquired SNPs, promoting the integration of longer DNA sequence. Despite its reliance on homologous recombination, we observed that transformation allows acquisition of highly polymorphic sequences (Fig. S6), likely facilitated by RecA tolerance for mismatches (*54*). Extended recombination events may maintain linkage between distant mutations and preserve combinations of alleles that may be beneficial together. Epistatic interactions are known to be important within proteins and between proteins of the same macromolecular complexes, which tend to be encoded in operons. A recent study showed that most operons in *E. coli* are less than 6 kb (*55*), suggesting that extended recombination provided by the transformation helicase/nuclease pairs may indeed facilitate the co-transfer of groups of SNPs in epistatic interaction, increasing the likelihood that genetic exchanges improve the recipient’s fitness. By bridging ssDNA invasions to form a continuous and extended recombination event, the helicase/nuclease system would allow the co-inheritance of linked variants, analogous to the maintenance of haplotypes.

## Materials and Methods

### Bacteria and growth conditions

All the *Legionella pneumophila* strains are derived from the Paris or Lp02 strains. They were cultured in solid medium containing ACES [N-(2-acetamido)-2-aminoethanesulfonic acid] and buffered charcoal yeast extract (CYE) or in liquid medium containing ACES and buffered yeast extract (AYE). When appropriate, media were supplemented with kanamycin (15 μg/mL for Paris strain and 50 μg/mL for Lp02 strains), streptomycin (50 μg/mL), gentamycin (10 μg/mL) or apramycin (15 μg/mL). All the *Acinetobacter* strains are derived from *A. nosocomialis* M2 or *A. baylyi* BD413 (also known as ADP1). When appropriate, they were grown in LB (Lennox formulation) broth or agar supplemented with tetracycline (5 μg/mL), kanamycin (50 μg/mL), apramycin (30 μg/mL), or rifampicin (100 μg/mL). The parental *Streptococcus pneumoniae* strain used in this study is R1818, an unencapsulated serotype 2 strain deficient for MMR and uncapable of CSP production (*56*). R1818 is derived from the R800 strain, a R6A derivative (strain genealogy detailed in (*57*)). Cells were cultured at 37°C on CAT-agar solid medium supplemented with 4% (vol/vol) horse blood or in Todd–Hewitt liquid medium (BD Diagnostic System) plus 0.5% yeast extract (THY) or C+Y medium. When appropriate, media were supplemented with rifampicin (2 μg/mL) or tetracycline (1µg/mL). The parent *Vibrio cholerae* strain used was E7946. When appropriate, LB Miller media was supplemented with rifampicin (100 µg/mL), erythromycin (10 µg/mL), zeocin (100 µg/mL), and/or trimethoprim (10 µg/mL). All strains and plasmids are listed in Table S3 and S4.

### Transformation assays

For transformation assays of *L. pneumophila*, the Paris parent strain (denoted wild-type, WT) is a constitutively transformable strain obtained by introducing a premature stop codon in the post-transcriptional repressor of competence *rocC* (*58*, *59*). Overnight streak of the desired *L. pneumophila* Paris strain was resuspended in 1 mL sterile water to an OD_600nm_ of 1 to obtain ∼1×10^9^ CFU/mL. A 10-μL portion of this suspension (∼1×10^7^ CFU) was spotted on CYE plate supplemented with 0.5 mM IPTG when appropriate. Then, 10 μL of transforming DNA (in Figure 1B, a 4 kb-long PCR product encompassing the *rpsL* mutation conferring resistance to streptomycin (*60*), or in Figure 4F a 4 kb-long PCR product of *lpp2553* region interrupted by an apramycin resistance gene) was spotted directly on the bacteria spot and left absorbed onto the agar. Plates were incubated at 37°C for 24 h. Each spot was then resuspended in 200 μL sterile water and ten-fold serial dilutions were plated on non-selective and selective medium. Plates were incubated at 37°C for 72 h and CFU counting was performed. Transformation assays of *A. baylyi* were conducted as follows: 70 μL of overnight culture in LB liquid medium at 30°C of the desired strain was used to inoculate 1 mL LB. After 1 h of incubation at 30°C with shaking, and 200 ng, 400 ng (for *kan-lacZ* Figure 4F) of tDNA was added and the cultures were reincubated at 30°C with shaking for 1 h to overnight and followed by DNAse I treatment to stop DNA uptake. Ten-fold serial dilutions were then plated on non-selective or selective medium. Plates were incubated at 30°C overnight and CFU counting was performed. Slight modifications to this method were done for the recombination event mapping, see below. For transformation assays of *S. pneumoniae,* pre-competent cultures were grown at 37 °C in C+Y medium to OD_550nm_ 0.1 and incubated with 50 ng/mL synthetic competence stimulating peptide (CSP) for 7 min at 37 °C to induce competence. DNA was then added at saturating concentration: 1 to 100 ng/mL for PCRs, 5 µg/mL for chromosomic DNA, and 1 µg/mL for the pLS1 plasmid. Cells were incubated 20 min at 30 °C, followed by phenotypic expression at 37 °C for 120 min. These cultures were plated on 10 mL CAT-agar 4% (vol/vol) horse blood, followed by selection using a 10 mL overlay of CAT-agar containing the selective antibiotic. Plates were incubated at 37°C overnight and CFU counting was performed. For transformation assays of *V. cholerae*, all strains contained P_tac_-*tfoX* and Δ*luxO* mutations, which bypass the native signals required for competence induction (*3*). Cells were growth to late-log by rolling at 30°C in LB supplemented with 20 mM MgCl_2_, 10 mM CaCl_2_, and 100 µM IPTG. Then, 7 µL of this culture was diluted into 350 µL of Instant Ocean medium (IO; 7 g/L Aquarium Systems) for each transformation reaction. Next, 5-10 ng of tDNA was added. Two tDNA products were used: (1) a 5 kb PCR product containing a Rif^R^ point mutation and (2) a 6 kb PCR product containing a ΔVC1807::Erm^R^ mutation. Control reactions were performed where no tDNA was added. Reactions were incubated at 30°C static overnight. Then, 1 mL of LB was added and reactions were shaken at 37°C for 3 hours to outgrow. Reactions were then plated for quantitative culture on LB supplemented with the appropriate antibiotic to select for transformants (rifampicin or erythromycin) as well as on plain LB to quantify total viable counts. Plates were then incubated overnight at 30 °C and CFU counting was performed. Transformation frequencies were calculated as the ratio of the number of CFU counted on selective medium (*i.e.*, the transformants) to the number of CFU counted on non-selective medium (*i.e.*, total population). All transformation assays were performed at least three times independently (on different days). Primers used to amplifiy each PCR fragment used as tDNA are listed in Table S5.

### Conjugation assay

For conjugation assays of *L. pneumophila*, the Phil-1 p45 Gent^R^ donor strain and the Lp02 recipient strain were streaked on CYE solid medium from the −80°C frozen stocks and incubated 72 h at 37°C. These cultures were restreaked on a new CYE plate and incubated overnight at 37°C to obtain exponentially growing cells. Co-culture was performed by mixing donor and recipient cells each to OD_600nm_ 0.1 (∼10^8^ cells/mL) in AYE broth (2 mL per replicate, final 1:1 mix is OD_600nm_ 0.2) followed by incubation for 24 h at 30°C. Tenfold serial dilutions of the co-culture were spotted on CYE plates containing streptomycin (selecting for the Lp02 recipient strain) and gentamicin (selecting for the acquisition of p45::*aacC1*) and incubated for 72 h at 37°C and CFU counting was performed. Conjugation frequency is the ratio of the number of CFU counted on selective medium divided by the number of CFU counted on nonselective medium (streptomycin only).

### Phylogenetic analyses

21,105 complete bacterial genomes present in RefSeq were downloaded in March 2021. The dataset was searched for the following hmm profiles (hmmsearch from hmmer v.3.3.2) (*61*): PF02021.20 for *yraN*, the co-occurrence of PF13541 (chlI domain), PF01078 (MgChelatase) and PF1335 (MgChelatase C domain) for *comM*, PF18073 (rubredoxin2 domain) and PF13481 (AAA_25 domain) for *radA*, PF00154.24 for *recA*, PF02481.18 for *dprA*, and PF06054 for *coiA*. A protein was considered present in the genome when the sequence score of the alignment was above the gathering threshold score cutoff (--cut_ga) and covered more than 50% of the hmm profile. The Genome Taxonomy Database (GTDB) (*62*) (March 2022, version r207) was used to pick representative strains for every bacterial species and build a bacterial phylogenetic tree. Among the 21,105 bacterial genomes of the dataset, 4,185 strains were represented in GTDB tree. This subsampling retained ca. 80% of the genera initially present. This subset of the GTDB tree obtained with the get_subtree_withtips function from castor R package was used to draw a tree. The phylum annotations of the strains were provided by GTDB. The *Bacillota* (formerly *Firmicutes*), *Desulfobacterota* and *Nitrospirota* phyla were each divided into several phyla by GTDB. They were grouped together to have a unique phylum for each of them and simplify the reading of the tree. Two proteins were considered co-localized in a genome when they are at less than 10 genes of distance. The statistical significance of the co-occurrence of genes in a genome was corrected for the phylogenetic structure of *Bacteria*. A phylogenetic logistic regression (phyloglm function from phylolm R package (*63*)) was used, to which we provided the GTDB phylogenetic tree. The data were fitted using the ‘logistic_MPLE” method (btol=20, boot=100). We ancestrally reconstructed the ancestral states of *yraN* and *coiA* in terms of presence-absence in each node of the bacterial phylogenetic tree. This was done using the Marginal Posterior Probabilities Approximation (MPPA) prediction method and the F81 evolutionary model of PastML v.1.9.34 (*64*).

### Protein expression and purification

Plasmids expressing CoiA in *E. coli* were obtained by amplification of *coiA* ORF from the genome of pneumococcal R800 strain by PCR with oligonucleotides harboring EagI and NotI restriction sites (Table S1), respectively. After EagI and NotI digestion, *coiA* ORF was ligated to pKHS vector (a pET28 derivative, Kan^R^) digested by NotI.. In order to produce C-terminal His-tagged YraN in *E. coli*, *yraN* from *L. pneumophila* was amplified by PCR and introduced in pKHS vector by InFusion methodology. The point mutations in *coiA* (E105A, D95A) and *yraN* (E50A) were introduced by site-directed mutagenesis using an adapted protocol from the Stratagene “QuickChange” with only one mutator primer carrying the modified sequence and with both the Pfu turbo polymerase (Stratagene) and the Taq ligase (New England Biolabs) in the PCR reaction. All the proteins were expressed from pKHS plasmids in *E. coli* BL21-Rosetta(DE3)-pLysS (Cm^R^) cells (Novagen) and were purified as soluble proteins following different procedures according to the type of protein. For CoiA and CoiA variants expression, cells were grown at 37°C in 250 mL LB medium with Cm (10 μg/ml) and Kan (50 μg/mL). For YraN and its variant, The expression clones were grown at 30°C in (400 mL) LB medium with Cm (10 μg/ml), Kan (50 μg/mL), and 0.2% glucose. When OD_600nm_ reached 0.7, protein expression was induced by adding 0.3 mM IPTG followed by growth at 23°C for 5 to 6 h in the case of CoiA and 0.5 mM IPTG followed by growth at 30°C for 2 h for YraN. Cell pellets were collected by centrifugation (6000 g; 20 min; 4°C) and further resuspended in 15 mL solution containing 50 mM Na-Phosphate pH 7.5, 400 mM NaCl, and 10 g/L lysozyme (for CoiA) or 30 mL solution containing 50 mM Na-Phosphate pH 7.5, 300 mM NaCl, 10 mM imidazole (for YraN) before breaking cells and DNA with sonifier (Vibracell). After clarification by centrifugation (15000 g; 30 min; 4°C in JA20), the soluble extracts of CoiA proteins were diluted to 250 mM NaCl with 20 mM Na-Phosphate pH 7.5 buffer and injected first to an Hydroxyapatite column (Bio-scale Mini CHT Type I, 5mL BioRad) equilibrated with 20 mM Na-Phosphate pH 7.5 and 200 mM NaCl using a FPLC (Äkta purifier-10, GE Healthcare). The YraN proteins were injected onto a His-Trap crude 1 mL column and further purified by FPLC using a solution of 50 mM Na-Phosphate pH 7.5, 300 mM NaCl, and an increasing gradient of imidazole. CoiA and variants were further eluted with a solution of 20 mM Na-Phosphate pH 7.5 and 400 mM of NaCl. The collected fractions of CoiA were diluted at 300 mM of NaCl before injection on a Hi-Trap Q HP 1 mL column to remove residual DNA before binding on Hi-Trap heparin HP 2 mL column (GE Healthcare) and elution with a NaCl gradient. The YraN eluted fractions were diluted 2.5 fold in 50 mM Na-Phosphate pH 7.5 to reach 120 mM of NaCl, and the same procedure as CoiA was followed using a Hi-Trap heparin HP 1 mL column (GE Healthcare). Eluted CoiA proteins were then separated on a Superdex 75 16/600 sizing column (GE healthcare) equilibrated with 20 mM Na-Phosphate pH 7.5, 200 mM NaCl. Proteins were finally purified and concentrated on Hi-Trap SP HP 2 mL column with a step at 300 mM NaCl. For YraN proteins, the column was equilibrated with 50 mM Na-Phosphate pH 7.5, 200 mM NaCl. Part of the purified proteins was stored at −80°C after addition of 10% glycerol, 1 mM TCEP for longer storage or at −20°C after addition of 50% glycerol. Another part was dialyzed against 50 mM Hepes pH 7.6, 200 mM NaCl, 0,5 mM TCEP, 10% glycerol using PD miniTraP 25 buffer exchange column for buffer comparative assays on activities. The purity and the concentrations of the purified proteins were analyzed by Bradford assay, SDS-PAGE and Coomassie staining analysis. The elution profiles of intact proteins and its respective variants were compared on analytical Superdex 75 10/300 sizing column (GE healthcare) (see Figure S3C and E). RecA was purified according Marie L. *et al.*, 2017 (*27*). DprA-His was purified according Quevillon-Cheruel S. *et al*., 2012 (*65*).

### DNA templates used for nuclease and binding assays

Depending on the experiments, oligonucleotides were ordered with a Cy3 fluorochrome at the 5’ or 3’ end (Eurogentec) or radiolabeled at their 5’ end under standard conditions. Briefly, 10 pmol of oligonucleotide to be labeled were incubated with 20 μCi of gamma-33P and T4 polynucleotide kinase (NEB, Biolabs) in a total volume of 10 μL of 1X PNK reaction buffer (NEB, Biolabs) for 1 hour at 37°C. After inactivating the kinase at 65°C for 10 minutes, free gamma-33P was removed using a G25 microspin column (GE-Healthcare). To construct ss-dsDNA hybrids, 3 pmol of radiolabeled primer was incubated with 5 pmol of cold complementary oligonucleotide in a total reaction volume of 10 μL containing 100 mM NaCl, 20 mM Tris-Cl pH 7.5, 10 mM MgCl2, and 1 mM DTT. After heat denaturation (3 min at 95°C), the hybridization mixture was placed in a boiling water bath (500 mL) until it reached approximately 40°C (almost 1h) to ensure proper hybridization of the oligonucleotides (see Figure S7 and Table S5).

### Nuclease binding assays

Increasing amounts of the purified protein of interest were mixed with the target DNA in 10 μL reaction buffer containing 50 mM Tris (or Hepes) pH7.5, 50 mM NaCl, 10 mM Mg(OAc)_2_, 0.3 mM MnCl_2_, 0.5 mM TCEP, 0.1 mg/mL BSA, 6% glycerol, and were incubated 30 min at 37°C. DNA concentration was 5 nM for the pUC18 supercoiled DNA, and 10 to 50 nM for the linear single/double DNA target, which are labelled with either Cy3 fluorochrome (Eurogentec) or radiolabelled at their 5’ extremity with T4 polynucleotide Kinase (NEB, Biolabs) and gamma ^32^P following standard conditions. The method used for each experiment is specified in the figure legends. To evaluate the protein binding efficiency, half of the reaction was directed loaded on 1% agarose / 1X TAE gel for plasmidic DNA, or on 5% acrylamide (29:1) / 0.3X TBE gels for short linear DNA substrates, and submitted to electrophoresis. For nuclease assays, quenching and DNA deproteinization was done by adding 0.15% SDS, 3 mM EDTA, 0.6 mg/mL proteinase K to the other half of the reaction for 10 min at 37°C before loading on gel. Incubation of 5 min at 95°C was added as a denaturation step for complex branched DNA targets (Figure 3A) to separate DNA strands before loading on 5% acrylamide / 0.3X TBE gel. This facilitated the visualization of the labelled ssDNA cleavage. Plasmidic DNA was visualized by UV detection after ethidium bromide staining (G-Box, Syngene). Short linear DNA structures were either detected with a phosphorimager (Fuji) for radiolabelled (^33^P) probes or with a fluor imager (Typhoon Trio, Fuji-GE-Healthcare) with an Abs/Em of 532/580 nm for fluorescent Cy3-labelled DNA. Mapping of cleavage site at nucleotide level was assessed on urea-denaturing sequencing gel. The quantification of nuclease activity annotated under the gel images correspond to the ratio between cleaved radioabelled DNA to total radiolabelled DNA. The analysis was done using the ImageJ software.

### D-loop nucleolysis assay

The D-Loop assay was performed using 10 nM of a 100-mer 5’Cy3-oligonucleotide homologous to pUC18 sequence, 150 nM of purified DprA^His^, and 300 nM of purified RecA. A first incubation first of 10 min at 37°C was performed in a 10 μL reaction solution containing 10 mM Tris-Cl pH7.5, 50 mM NaCl, 10 mM Mg(OAc)_2_, 0.3 mM MnCl_2_, 2 mM dATP, 0.5 mM TCEP, 0.1 mg/mL BSA, and 6% glycerol to promote RecA loading on ssDNA (RecA nucleofilament). Then, 5 nM of the DNA target (supercoiled pUC18 vector) was added and further incubated for 10 min at 37°C to allow homologous pairing (D-Loop) between ssDNA and dsDNA. Increasing amounts of CoiA (See figure legend) was then added and incubated 30 min at 37°C. A negative control (no D-Loop) was performed in parallel with a non-homologous oligonucleotide. The reaction was stopped by addition of SDS/EDTA. After standard DNA extraction by chloroform and ethanol/salt DNA precipitation, the DNA was resuspended in 10 μL of water. A third of the extracted DNA was loaded for migration on 1.3% agarose / 1X TAE gel to evaluated D-Loop efficiency. Another third of the DNA was used as matrice for the primer extension experiment.

### Primer extension experiment

A total of 5 nM of radiolabelled (−40) primer complementary to the pUC18 vector was incubated with 1.5 nM of DNA matrix, 0.5 mM of dNTP mix, and the Thermosequenase ^TM^ DNA polymerase in the corresponding reaction buffer following the instructions (Thermosequenase^TM^ Cycle Sequencing kit, Fisher Scientific). Primer extension was performed by PCR for 10 three step-cycles of 30 sec at 95°C, 30 sec at 55°C and 30 sec at 72°C. In parallel, standard chain-termination DNA sequencing was performed on pUC18 with the same primer and ddNTP using the same kit for 50 PCR cycles, following the manufacturer instructions. Sequencing and primer extension reactions were loaded on 8 M urea – 6% acrylamide (19:1) – 1X TBE denaturing gel. After migration at 40 watts for 1.5 h, the gel was transferred to 3 MM Whatmann paper, dried, and exposed overnight for phosphorimager analysis.

### Bacterial two-hybrid system experiments

Two hybrid experiments were carried out to test the interaction between proteins involved in transformation-associated homologous recombination. To that end, *recA*, *dprA*, *yraN* and *comM* genes from *L. pneumophila* Paris were cloned into pKNT25 and pUT18 plasmids in frame with T25 or T18 catalytic domains of *B. pertussis* adenylate cyclase (*47*) by Sequence and Ligation Independent Cloning (SLIC) (*66*). With this method, for each protein of interest, two recombinant plasmids encoding two hybrid proteins (with the protein of interest in N-ter) were propagated in their respective *Escherichia coli* DH5α competent hosts: pKNT25-X and pUT18-X, encoding respectively X-T25 and X-T18. Each plasmid encoding an X-T25 fusion protein was then co-transformed in competent BTH101 *cya*^-^ strains with a plasmid encoding X-T18 fusion protein. The co-transformed bacteria obtained were pre-cultured in LB liquid medium supplemented with 25 μg/mL kanamycin and 100 μg/mL ampicillin at 37°C overnight with shaking. After incubation, these cultures were 1:10 diluted and spotted onto LB agar supplemented with 25 μg/mL kanamycin, 100 μg/mL ampicillin, 0.5 mM IPTG and 100 μg/mL X-β-Gal (Roth) and incubated at 20°C for 48 to 72 h. Interaction between the two RecA constructs are used as positive controls in this experiment since its interaction was already described (*48*).

### Comigration marker experiment

Comigration marker experiment was adapted from the comigration assay conducted in *V. cholerae* (*30*). This experiment was conducted in a *A. nosocomialis* M2 strains that carry genetically linked markers: the *lacZ* gene placed adjacent to a tetracycline cassette. This strain called “lacZ+-tet” was naturally transformed with the following PCR products: “2kB-lacZ^dlt1^-kan-2kB” or “2kB-lacZ^dlt2^-kan-2kB” or “2kB-lacZ^dlt3^-kan-2kB” as tDNA. These tDNA have 2 kB upstream and downstream homologous regions with “lacZ+-tet” recipient strains that allow the acquisition of the tetracycline cassette (replacing the kanamycin cassette) and of the 7-bp deletion in *lacZ* (lacZ^dltX^) by homologous recombination. The 7-bp deletions were designed so that (i) they are not substrates for the mismatch repair system and (ii) they shift the reading frame of *lacZ* to produce a TGA stop codon resulting in *lacZ*^-^ phenotype. These 7-bp deletions were designed at increasing distances from the original TAA stop codon of the kanamycin cassette (distance between kan_TAA_ and lacZ_TGA_ for dlt1=253 bp, for dlt2=1,606 bp, and for dlt3=3,043 bp).Transformation assays of *A. nosocomialis* were performed as previously described (*67*, *68*). 2 mL LB were inoculated with overnight streak of the desired strain and incubated at 37°C until reaching an OD_600nm_ of 1 to obtain ∼1×10^9^ CFU/mL. The culture was 100-fold diluted in sterile water to obtain 10^7^ CFU/mL. For each mutant, 5 μL of this suspension was mixed with 5 μL of purified PCR (150 ng/μL) dlt1, dlt2, or dlt3, followed by incubation at 37°C for 24 h. Ten-fold serial dilutions of the transformation cultures were plated on LB agar supplemented with 50 μg/mL kanamycin and 100 μg/mL X-gal or on nonselective LB plates and incubated in appropriate conditions described before. Comigration efficiency, which refers to the number of kanamycin-resistant transformants that have acquired the *lacZ* deletion, was calculated as the ratio of the number of white transformants (Kan^R^/*lacZ*^-^) divided by the number of total transformants (Kan^R^/*lacZ*^+^ + Kan^R^/*lacZ*^-^) for each tDNA and each mutant.

### Characterization of recombination events

Direct detection of transformation recombination events was carried out as follows. *A. baylyi mutS* null and *S. pneumoniae* strains were transformed with a 6.4 kb-long PCR product as homeologous tDNA that confers resistance to rifampicin upon recombination. This tDNA was obtained by amplification of *A. nosocomialis* M2 rifR genome (*69*) or *S. mitis* NCTC 12261 [NS 51] (*70*) rifR genome that carries the *rpoB* gene, with the selected rifR mutation (Q522L A1565T in M2, G5079A (n=194) and G5079T (n=2) in *S. mitis*) at a distance of 1.4 kb from the closest fragment extremity (see primer list in the Table S5). Of note, 68 out 263 transformants in *Sp* which displayed recombination events were kept despite uncertainties on the identity of the acquired rifR mutation. Parental and mutant *A. baylyi* strains were transformed with 1 µg/mL of DNA for 6 h and plated on rifampicin-containing LB plates. Parental and mutant *S. pneumoniae* strains were transformed as described above with 100 ng/mL of DNA. In each of three independent experiments, 24 to 32 colonies were picked for each tested strain and the region of interest was amplified as PCR products. In the case of *A. baylyi mutS* strains, a detection PCR was performed to differenciate transformants from spontaneous rifR mutants before sequencing (see Table S5). The validated 6.4 kb-long PCR fragments were used to construct a sequencing library (Rapid Barcoding Kit 96 SQK-RBK110.96, Oxford Nanopore) and sequenced using a MinIon instrument (Oxford Nanopore). For *Sp*, sequencing was performed using R9.4 flow cells and reads were basecalled with the Guppy 6.1.3 using super accurate dna_r9.4.1_450bps_sup model and filtered on coverage. Because the donor population was heterogeneous (194 with G5079A mutation, 2 with G5079T mutation, and 68 samples without identified rifR determinant), a theoretical *S. mitis* sequence was used. Variant calling was performed using Medaka (1.5.0) with medaka_haploid_variant using the donor and recipient sequence as reference. Quality score of called positions was checked, all variable positions were kept (score > 2). Sequencing depth was checked with bedgraphs obtained based on alignments using samtools (1.10) and bedtools (2.27.1), and then visually checked using ggplot2. Samples displaying low sequencing depth were removed. All samples displayed recombination event, therefore they were all kept. For *Ab*, sequencing was performed with R10.4.1 flow cells and basecalling was performed using Dorado (7.2.13) with super-accurate model. Reads were aligned on the reference sequence (*A. nosocomialis* donor) with minimap2 (2.28-r1209) and indexing of alignment was done with samtools. Clair3 (1.0.10) was used for variant calling with haploid_precise option and r1041_e82_400bps_sup_v500 model. Five variable positions were removed from further analyses due to low variant calling score (tag LowQual in VCF) and/or multiple substitutions inconsistent between samples. Sequencing depth was checked with bedgraphs obtained based on alignments using samtools (1.20) and bedtools (2.30.0), and visually checked using ggplot2 (3.5.1). PCR success was confirmed for all samples. Presence of the rifR A1368T mutation was checked in samples’ VCF. Spontaneous mutants (samples not carrying rifR A1368T) were removed. For both species, Samples VCF were compared to recipient on donor reference VCF. A position was considered recombinated in the sample if it was absent in sample’s VCF (site is identical to the donor) and present in reference’s VCF (site is a variable site between donor and recipient).

### Statistical analysis

Normality of distributions was visually inspected using qqplots. Homoscedasticity between samples was checked using Levene’s test. Events length means were compared using Welch’s ANOVA, and post-hoc pair comparisons were conducted with Games-Howell test. For SNP and segment number and mean segment length, non-parametric analyses were performed using Kruskall-Wallis test. Post-hoc pair comparisons were performed using Dunn’s test. All statistical tests were done with R package rstatix. Distributions plots were build using ggplot2, PupillometryR and cowplot packages.

### Construction of mutants of genes involved in natural transformation

For mutation of *Acinetobacter* and *Legionella*, the target ORF was interrupted by a resistance cassette in the case of gene inactivation, or replaced by a mutant allele in the case of point mutations. To this end, the ∼2 kB upstream region (PCR product A [PCRA]) and the ∼2 kB downstream region (PCRC) were amplified by PCR respectively using primer pairs S_X_P1/S_X_P2-tail-Y and S_X_P3-tail-Y/S_X_P4 (where S designates the strain, X the targeted gene and Y the type of resistance cassette). X_P2-tail-Y and X_P3-tail-Y carry ∼30-nucleotide sequences complementary to the ends of the Y cassette. This complementarity was used to assemble the PCRA and PCRC with the Y cassette (PCRB, amplified with Y_F/Y_R primer pair) by overlap extension PCR. Overlapping PCR products were naturally transformed in appropriate conditions for each organism and plated on the corresponding selective media. The *Vibrio* mutant strains were obtained using the MuGENT method (*71*, *72*). In *Sp*, null mutation of coiA was performed as previously described (*73*). Briefly, plasmid pR410 was used as the source of the 1,337 bp mariner (Kan^R^) mini-transposon. Plasmid DNA (∼1 µg) was incubated with *coiA* PCR fragments (∼1 µg with coiA1 and coiA2 oligonucleotides) in the presence of purified Himar1 transposase (*74*) in a total volume of 40 µL, resulting in the random insertion of the mini-transposon within the fragments. Gaps in the transposition products were repaired as previously described (*75*), and the resulting in vitro-generated transposon insertion library was used to transform S. pneumoniae. Kan^R^ mutants were characterized by sequencing. Point mutations in *coiA* were introduced at the native *coiA* locus in R1818 by natural transformation with pKHS-coiA^D95A^ and pKHS-coiA^E105A^ without selection, and clones were screened by sequencing the *coiA* locus. The primers used for the strain’s construction are listed in Table S5.

### Construction of mutants for comigration marker experiment

Mutants for the comigration marker experiment were constructed as follows. A first construct was generated as described above with few modifications. First, a cassette (PCRB) comprising the *lacZ* gene upstream of the tetracycline resistance cassette which are antisense to each other (“*lacZ-tet*” cassette) was generated by assembly PCR of the *tetA* and *lacZ* genes using strgPRM_lacZ_F forward primer (that allows the addition of a strong promoter upstream of *lacZ*) and tcR-tail-attTn7_F reverse primer. Then, 2 kB upstream (PCRA: 2kB-attTn7_P1/attTn7-P2) and downstream (PCRC: TcR-2kB_P3/2kB-attTn7_P4) regions homologous to the integration region of the attTn7 site were added to the PCRB “ *lacZ-tet*” cassette by assembly PCR. This “2kB-*lacZ-tet*-2kB” construct was directly used to transform *A. nosocomialis* strains M2 WT, M2 *yraN*, M2 *comM* and M2 *yraN comM* to form the different “*lacZ-tet*” recipient strains (genotype *attTn7*::*lacZ-tetA*) of this experiment. The “*lacZ-ka*n” strain was generated in the same way only in the wild-type M2 background. DNA from this strain was used as a template to generate the *lacZ* deletion constructs (“*lacZ^dltX^-kan*” with genotype *attTn7*:: *lacZ^dltX^-aph*). A first PCR (PCRA) was performed with the attTn7-2kB_P4 primer and the lacZ-dltX-tailed_P3 primer which was designed to generate a 7-bp deletion in *lacZ*. This primer also has a complementary tail to the end of the PCRB fragment. The PCRB fragment is generated with the primers 2kB-attTn7_P1 and lacZ-dltX_P2 which is complementary to the tail of the lacZ-dltX-tailed_P3 primer. PCRA and B were assembled by overlap extension PCR using 2kB-attTn7_P1 and attTn7-2kB_P4 to finally generate the following construction: 2kB-*lacZ^dltX^-kan*-2kB which is used to transform “*lacZ-tet*” strains. Finally, the DNAs from these strains were used as a template to generate the tDNA used for the comigration marker experiment (2kB-*lacZ^dlt12,3^-kan*-2kB) by PCR with 2kB-attTn7_P1 and attTn7-2kB_P4 primers.

### Strain construction for bacterial two-hybrid experiments

Plasmids pUT18 and pKNT25 were first linearized using the primers pUT18_F/R or pKNT25_F/R, respectively. The genes to be cloned into these vectors are amplified by PCR with X-pUT18_F/R (for cloning into pUT18) and X-pKNT25_F/R (for cloning into pKNT25) which allowed addition of complementary regions to the ends of the linearized vector. All linearized vectors and inserts are purified on gel and treated with DpnI. The SLIC reaction is then performed in a final volume of 20 µL, completed with water by adding a 3:1 ratio of insert:vector with 0.5 µL of T4 DNA polymerase and 2 µL of NEB1.2 buffer. This mix is incubated at room temperature for 2 minutes and 30 seconds. The product of the SLIC reaction (which theoretically corresponds to the recirculated vector with the insert) is directly used for cloning into *E. coli* DH5α competent cells.

### Protein immunodetection

Cultures for immunodetection of CoiA and Ssb proteins were performed as described previously (*27*). Briefly, 30 mL of each pre-competent *Sp* culture (*coiA* WT, *coiA*^D95A^ and *coiA*^E105A^) grown at 37°C in CH medium to OD_550_ 0.1 were incubated at 37°C with 25 ng.mL^-1^ synthetic competence stimulating peptide (CSP) to induce competence during 0 to 60 min. After centrifugation 5 min at 5,000g at 4°C, cell pellets were stored at −80°C. Pellets were latter adjusted at OU∼2 with lysis buffer (TE 1X and 0.01% DOC). A fraction (1/5) of each sample was supplemented with Laemmli SDS-PAGE loading buffer, heated for 5 min at 80°C and loaded on SDS-PAGE precast 4–20% gels (Mini-PROTEAN TGX gel, Bio-Rad). After electrophoresis and protein transfer to nitrocellulose membrane (Mini Nitrocellulose Transfer Packs, Bio-Rad) with the trans-Blot Turbo Transfer System (Bio-Rad), membrane was cut above the 17 kDa marker band. The top part of the membrane was probed with anti-CoiA antibodies (1:1,000) and the bottom part with anti-SsbA and anti-SsbB mix (1:5,000 each). All antibodies were raised in rabbit (Eurogentec). The YraN-FLAG produced in *Ab* were detected using a commercial kit (Sigma) following the manufacturer instructions.

## Supporting information

Supplementary materia

## Acknowledgments

Acknowledgments follow the references and notes list but are not numbered. Start with text that acknowledges non-author contributions (including disclosure of any editing services or AI-assisted technologies used in preparation of the manuscript), then complete each of the sections below as separate paragraphs.

## Funding

Agence Nationale de la Recherche ANR-20-CE12-0004 (XC, EPCR)

Agence Nationale de la Recherche ANR-10-BLAN-1331 (PP)

Agence Nationale de la Recherche ANR-17-CE13-0031 (PP)

Agence Nationale de la Recherche ANR-22-CE44-0044 (PP)

Computational and storage services (TARS cluster) provided by the IT department at Institut Pasteur, Paris

Agence Nationale de la Recherche ANR-10-LABX-62-IBEID (EPCR)

Agence Nationale de la Recherche PIA/ANR-16-CONV-0005 (EPCR)

Fondation pour la Recherche Médicale FDT202001010890 (LH)

European Union’s Horizon research and innovation programme. Marie

Skłodowska-Curie Postdoctoral Fellowships grant 101208987 (CJR)

National Institutes of Health R35GM128674 (ABD).

## Author contributions

XC and PP conceptualized the project; LH performed genetic analyses and transformation assays on Lp and Ab; VM conducted all biochemistry analyses; MB performed genetic analyses and transformation assays on Sp; TND and ABD performed genetic analyses and transformation assays on Vc; FM and EPCR performed phylogenetic anayses; LH, MB and CJR acquired data on recombination events; JP analysed recombination events analysis; CJR consolidated data, prepared figures and drafted the manuscript; All authors contributed to final edition of the manuscript.

## Data and materials availability

All data are available in the main text or the supplementary materials.

## Supplementary Materials

Table S1. Phylogenetic distribution.

Tables S2–S5

Figs. S1 to S7

