## Supplementary materia for "Two D-loop resolution systems enable natural genetic transformation in bacteria"

**The PDF file includes:**

Figs. S1 to S7
Tables S2 to S5

**Other Supplementary Materials for this manuscript include the following:**

Table S1

### Figure S1

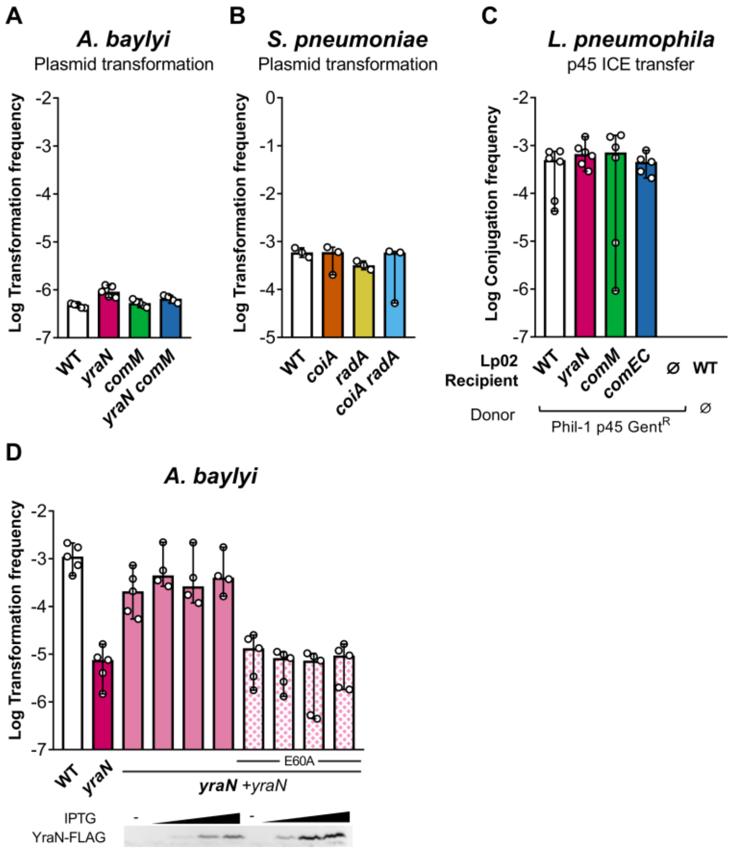


**Fig. S1. Natural transformation of plasmid and conjugation of ICE.** (A) Transformation frequencies of *A. baylyi* (*Ab*) and mutants by the pX5K plasmid capable of episomal replication in a wide range of hosts and carrying the kanamycin resistance gene. *Ab* was exposed to the purified plasmid for 24 h at 30°C and cultures were plated on selective plates containing kanamycin. Transformation frequencies represent the ratio of colony-forming units on selective over the total CFU count on non-selective plates. (B) Transformation frequencies *S. pneumoniae* (*Sp*) and mutants by the broad host range streptococcal plasmid pLS1, capable of episomal replication and carrying the tetracycline resistance gene. *Sp* was exposed to the purified plasmid for 20 min at 30°C. Transformation frequencies represent the ratio of colony-forming units on selective over the total CFU count on non-selective plates. (C) Conjugation efficiency of the p45 integrative conjugative element (ICE) carrying a gentamicin resistance gene (Gent$^R$) from the *L. pneumophila* Philadelphia-1 strain to the parental *L. pneumophila* Lp02 strain (WT) or *yraN*, *comM*, and *comEC* mutants. The latter demonstrates that transfer of p45 is indeed independent of natural transformation. Control experiments were performed without recipient or without donor strains. Following co-culture in AYE broth for 24 h at 30°C, cultures were plated on selective and non-selective CYE plates. Conjugation frequencies represent the ratio of colony-forming units on selective over the total CFU count on non-selective plates. (D) Controls related to Fig. 1D. Transformation frequency of the *yraN-flag* allele-carrying complementation strain and immunodetection of the YraN-FLAG, tested at IPTG concentrations of 0, 0.1, 0.5 and 1 mM.

### Figure S2

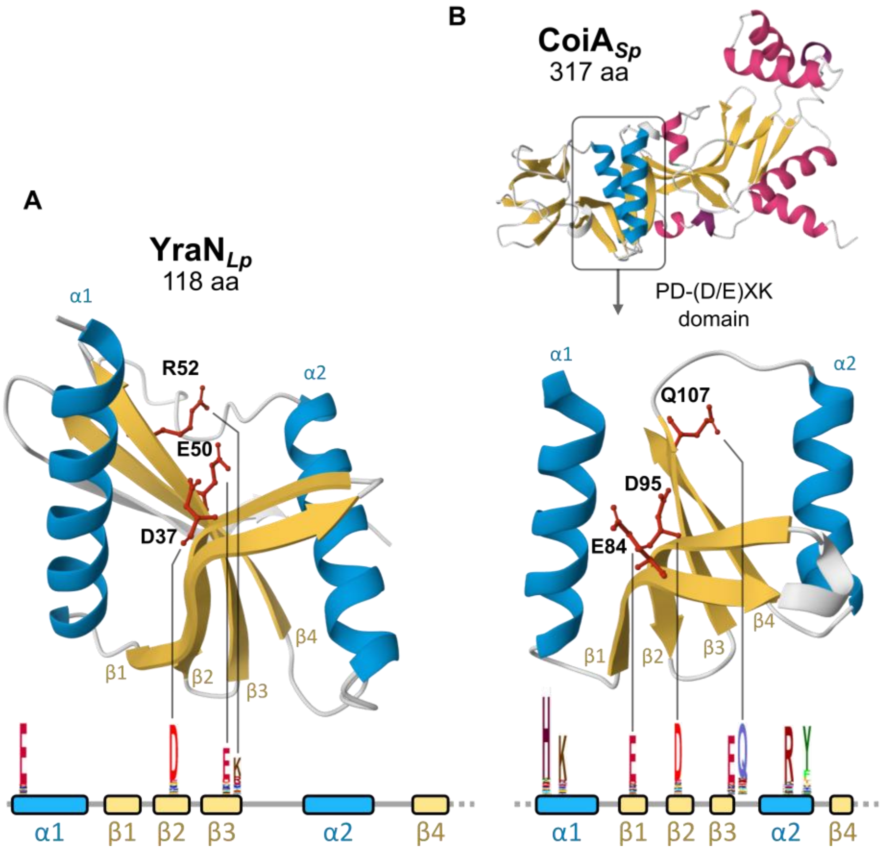


**Fig. S2. CoiA and YraN are predicted PD-(D/E)XK phosphodiesterases.** Predicted structure of the *L. pneumophila* YraN (A) and *S. pneumoniae* CoiA (B) using AlphaFold 3. Full CoiA (top) and its nuclease domain are depicted. Both proteins display the typical fold of PD-(D/E)XK phosphodiesterase consisting of four beta strands (yellow) sandwiched by two alpha helices (blue). Alpha helices outside of the nuclease domain of CoiA are depicted in pink. The linear representation show the highly conserved residues indicated as weblogo of the UPF0102 (A) and PF06054 (B) domains. Some conserved residues in the catalytic sites are highlighted in the 3D structures.

#### Figure S3

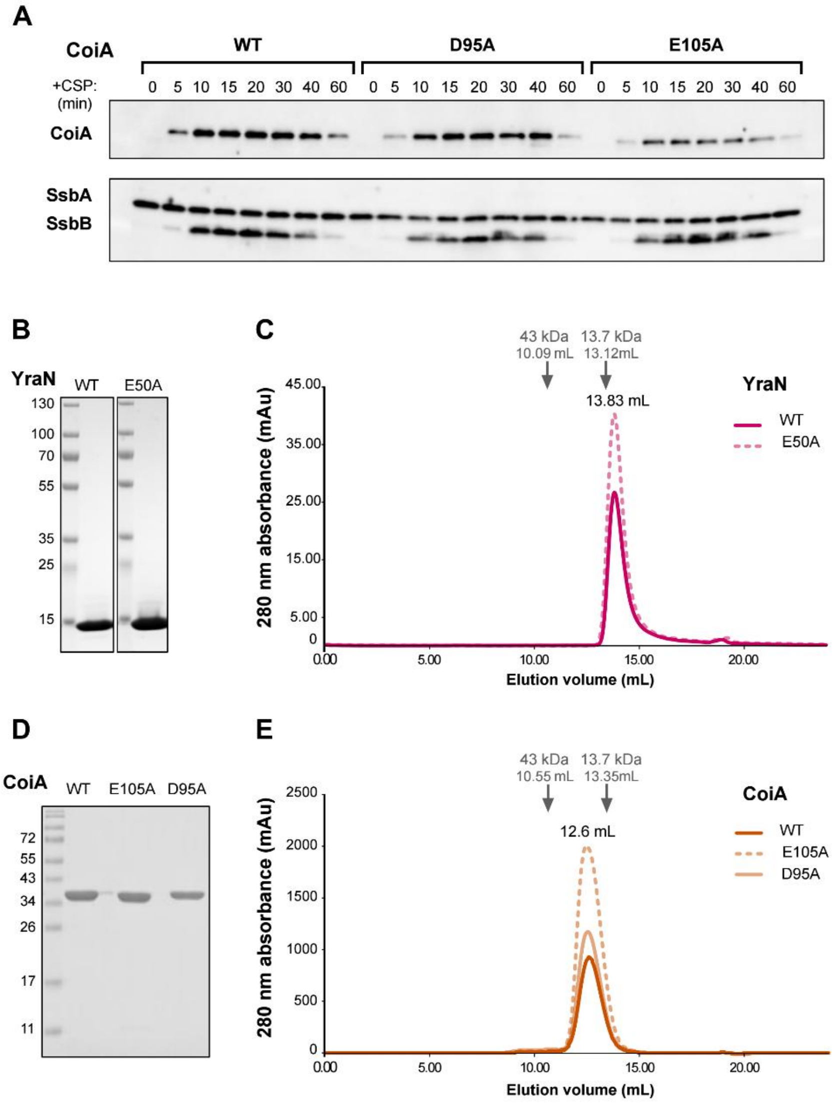


**Fig. S3. Expression and elution profiles of CoiA and YraN$_{Lp}$.** (A) Expression in S. pneumonia of wild-type CoiA and mutated derivatives (E105A and D95A), at different time after CSP induction, detected by immunoblot and CoiA specific antibodies. Co-detections of inducible SsbB and constitutive SsbA are performed in parallel for CSP induction control and sample standardization, respectively. (B) Profile of C-terminal His tagged YraN$_{Lp}$ and E50A mutant proteins expressed in *E. coli* and purified to homogeneity, migrated on SDS-PAGE with Coomassie staining. (C) Elution profile of wild-type YraN and E50A mutant on analytical SEC (Superdex 75 PG 10/300). Ribonuclease A (13.7 kDa) and Ovalbumine (43 kDa) were used as size markers and their elution profiles are indicated with arrows. (D) Profile of purified wild-type CoiA and catalytic site mutated derivatives E105A and D95A after migration on SDS-PAGE and Coomassie staining. E. (as in C) Elution profile of CoiA and variants on analytical SEC (Superdex75 10/300).

### Figure S4

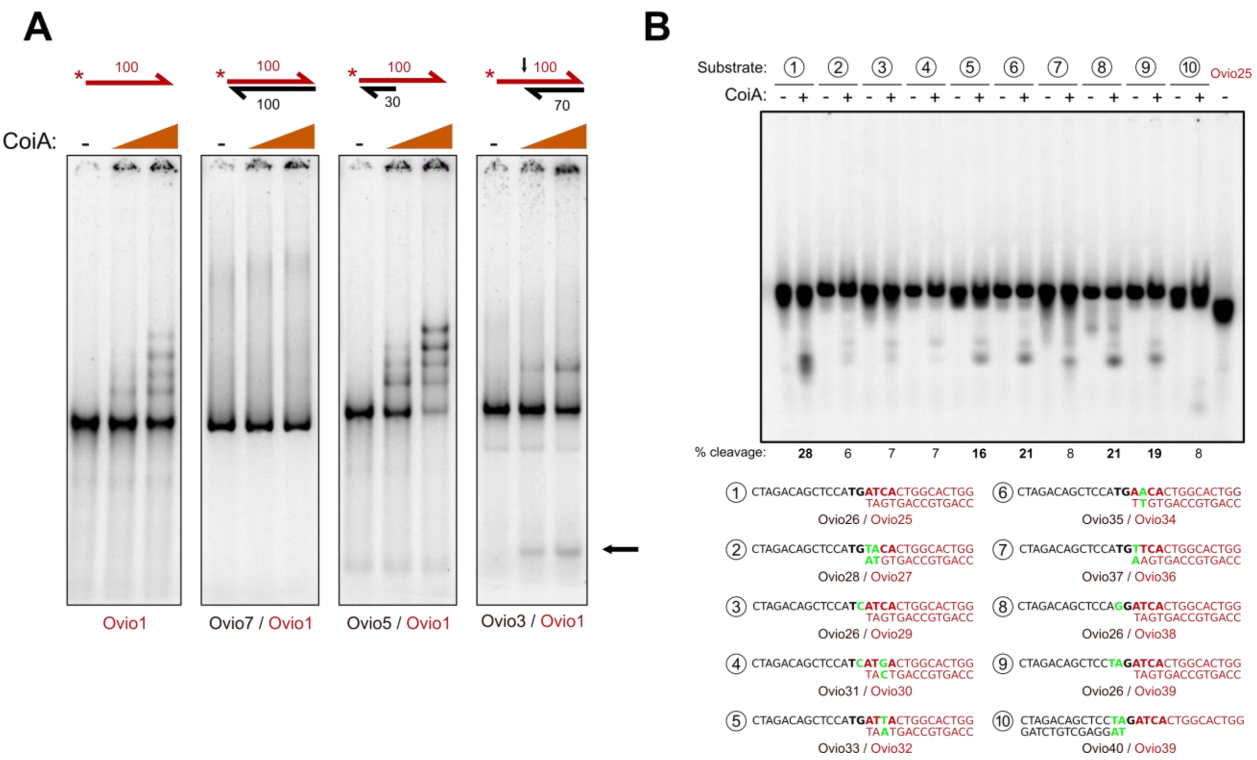


**Fig. S4. CoiA interaction generates a ladder-like shift pattern and is more stable with ssDNA than with dsDNA.** (A) Binding assays performed on native PAGE with increasing amounts of CoiA (30; 100 nM) tested on various DNA substrates, schematized on top of each panel. The numbers indicate the size in nucleotides of the oligonucleotides. Half arrow extremities symbolize 3'-ends of DNA strands. The black arrow points to the CoiA cleavage site. The 5'-radiolabelled (*) oligonucleotide is depicted in red. The black arrow next to the gel shows the band resulting from substrate cleavage by CoiA. Oligonucleotides used to construct the structures are detailed below the gels. (B) CoiA activity on ten 5'-tailed DNA structures with various ss-dsDNA junction sequences. Top: nuclease assays visualized on PAGE after heat denaturation and protein removal. Bottom: sequences of the ten DNA templates.

### Figure S5

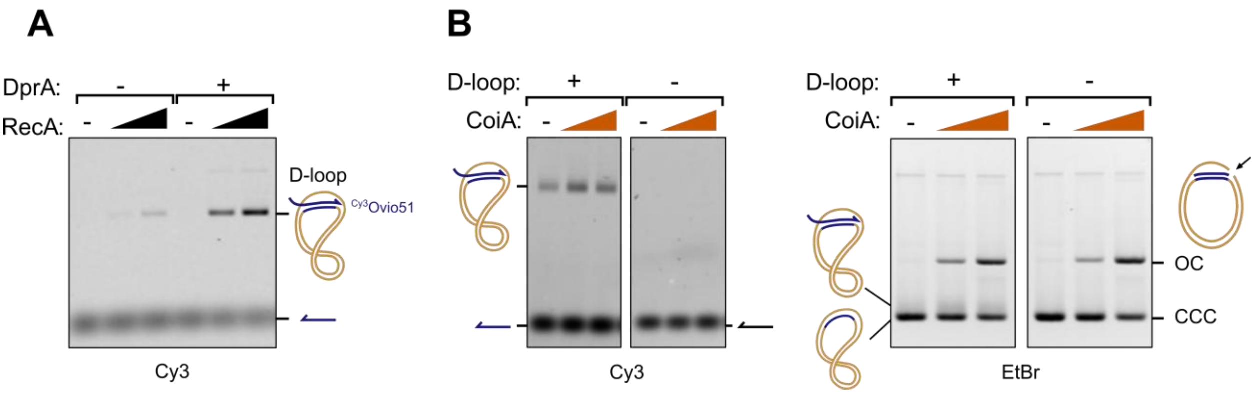


**Fig. S5. D-loop nucleolysis assay design and validation.** (A) D-Loop assay optimization with RecA and DprA. Hybridization of a 100-mer fluorescent (Cy3) oligonucleotide ($^{Cy3}$Ovio51, blue) with its complementary sequence on supercoiled ds-DNA was assessed without or with RecA (150; 300 nM) and DprA (150 nM) *Sp* proteins. The efficiency of D-Loop was analyzed after removal of the proteins, separation of the DNA molecules on agarose gel, and detection of the Cy3 fluorescent signal. (B) D-Loop formation controls as CoiA nuclease activity associated to Fig. 3C-D. Analysis of hybridization of a 100-mer fluorescent oligonucleotide (blue arrow) with a complementary sequence on supercoiled ds-DNA or CCC DNA (left panels, + D-Loop) in presence of DprA (150 nM) and RecA (300 nM), with or without CoiA (130; 400 nM). DNA molecules were separated on agarose gel after protein removal. The detection of fluorescent signal (Cy3) determined the D-Loop proportion whereas the Ethidium Bromide staining (EtBr) of the same gel revealed the efficiency of cleavage of CCC DNA by CoiA. As a negative control for D-Loop, same experiment was realized with a fluorescent ss-DNA that has no homology with the ds-DNA (right panels, - D-Loop). The samples were further used for primer extension experiment shown in Fig. 3D. BET (right). “CCC”: Covalently Closed Circle, “OC”: Open Circle.

### Figure S6

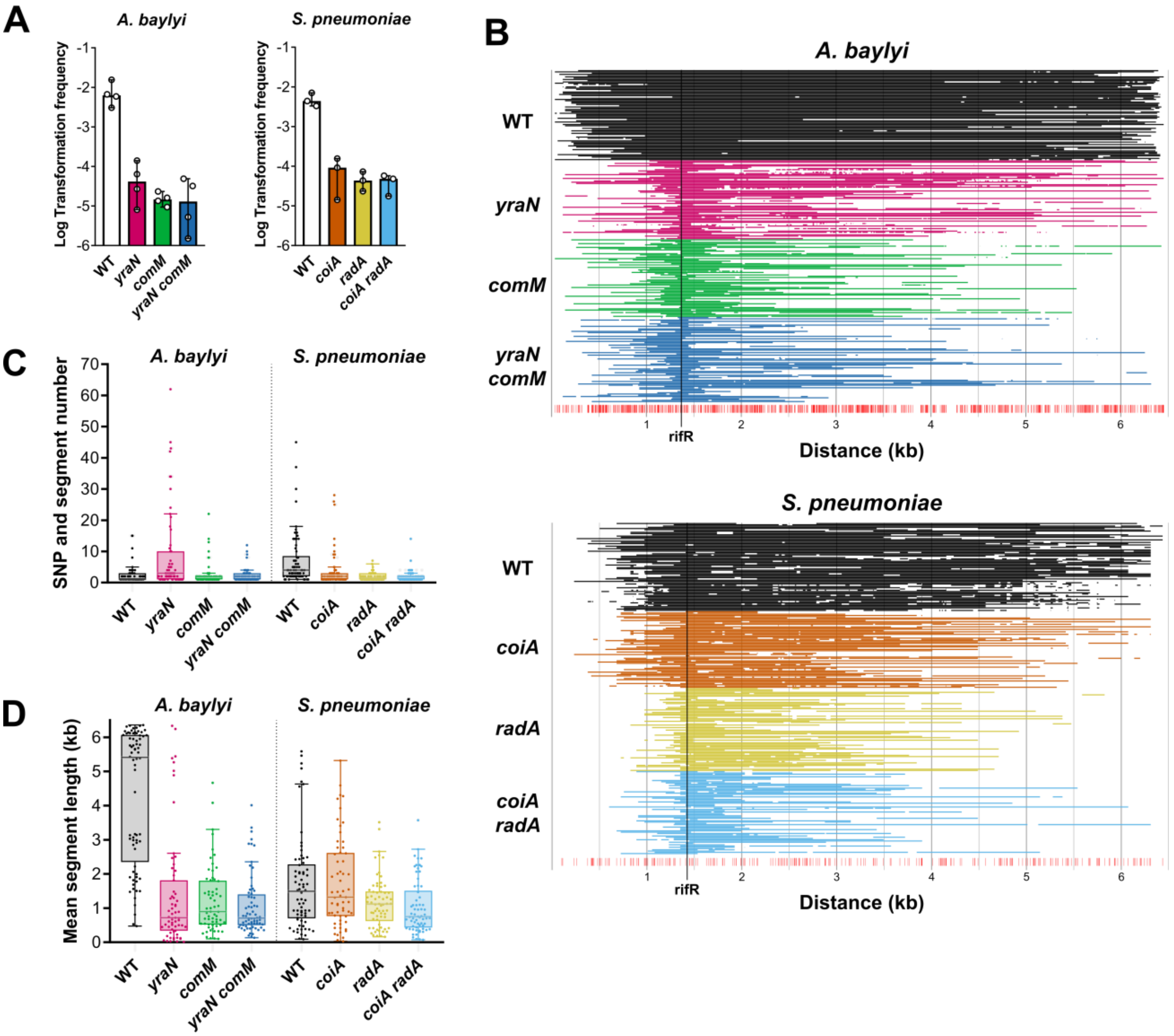


**Fig. S6. High-resolution mapping of recombination events.** Data from the recombination events mapping assay illustrated in Fig. 5A: Parental and mutant strains of *Ab* (*mutS* null) and *Sp* were naturally transformed with a 6.4 kb-long homeologous PCR fragment carrying an off-centered a $\mathrm{Rif}^{R}$-conferring SNP. PCR fragments were amplified from *A. nosocomialis* M2 (*An*, carrying 481 SNPs relative to the *Ab* recipient region), and from *S. mitis* (*Sm*, carrying 251 SNPs relative to the *Sp* recipient region). The target region was sequenced in 62 to 72 transformants for each strain. (A) Efficiency of natural transformation. Transformation frequencies represent the ratio of colony-forming units on selective (containing rifampicin) over the total CFU count on non-selective plates. (B) Reconstruction of recombination events on the basis of the acquired SNPs. Individuals SNPs (bottom red lines) distinguishing the *An* and *Sm* donors DNA and the *Ab* and *Sp* recipients were identified in the sequences of the transformants. Each line corresponds to an individual transformant. Two or more successive acquired SNPs define a recombination segment (displayed as lines). Individual SNPs that are flanked by non-acquired SNPs are also displayed, however can be difficult to see at this scale. The position "rifR" corresponds to the SNP conferring resistance to rifampicin which was used to select transformants. (C) Total number of acquired segments and individual SNPs (see Fig. 5A for metrics definition) for each transformant. (D) Mean length of acquired segments of each transformant, defined as regions of at least two successive converted SNPs. Whiskers delineate the range, boxes delineate the 1st and 3rd quartiles, horizontal line shows the median.

### Figure S7

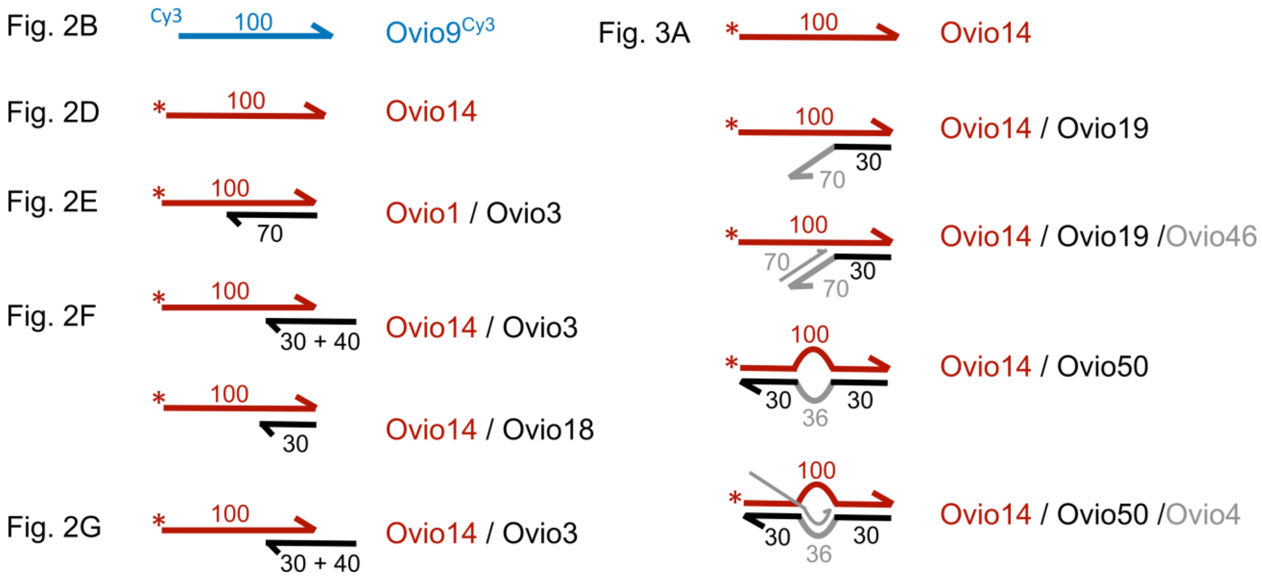


**Fig. S7. Oligonucleotides used to construct the target DNA structures.** Schematic representation of the various DNA substrates used in this study. The oligonucleotides sequences are listed in Table S5.

**Table S2**. Genes required for competence and natural transformation in both *L. pneumophila* and *S. elongatus*.

Data were retrieved from previous studies (Hardy et al., 2021; Taton et al., 2020). CDS were consistently annotated using eggNOG-mapper (Huerta-Cepas et al., 2017) and orthologous genes in *L. pneumophila* and *S. elongatus* were matched based on gene name and/or COG when available. Values correspond to the $\log_2$ fold change ($\log_2$FC) in abundance of insertions in the corresponding genes between the population obtained following transformation and the control, (non-transformed, population). Data were sorted and filtered for $\log_2FC < -1$ and with P-value < 0.05.

| Gene name | COG | Annotation | **Ratio transformed vs control** (log2 fold change) | |
|---|---|---|---|---|
| | | | ***L. pneumophila*** | ***S. elongatus*** |
| *comM* | COG0606 | ComM DNA helicase | -3.3 | -1.4 |
| *comEC* | COG0658 | DNA transport transmembrane protein ComEC | -2.9 | -2.4 |
| *comEA* | COG1555 | Competence protein ComEA | -3.5 | -2.5 |
| *pilM* | COG4972 | Tfp pilus assembly protein PilM | -3.0 | -2.9 |
| *pilN* | COG3166 | Tfp pilus assembly protein PilN | -3.6 | -2.8 |
| *pilO* | COG3167 | Tfp pilus assembly protein PilO | -2.7 | -2.6 |
| *pilP* | COG3168 | Tfp pilus assembly protein PilP | -2.6 | -2.4 |
| *pilQ* | COG4796 | Tfp pilus assembly protein PilQ | -3.6 | -2.7 |
| *pilB* | COG2804 | Tfp pilus assembly protein PilB | -3.7 | -3.1 |
| *pilC* | COG1459 | Tfp pilus assembly protein PilC | -3.3 | -3.1 |
| *comFC* | COG1040 | Competence protein ComFC | -3.0 | -2.0 |
| *dprA* | COG0758 | DNA protection and recombination DprA | -2.7 | -2.3 |
| *yraN* | COG0792 | UPF0102 protein, unknown function | -2.8 | -1.0 |

**Table S3**. Strains used in this study

| Strain name in manuscript | Genotype, antibiotic resistance, use | Reference & strain number |
|---|---|---|
| | ***Streptococcus pneumoniae*** | |
| WT (*Sp*) | *comC0 hexAΔ3*::*ermAM (*Ery$^{R}$), also labeled as R1818<br>Denoted as wild-type in this study. | (Attaiech et al., 2011)<br>R1818<br>*Strain genealogy detailed in*<br>(Prudhomme et al., 2016) |
| rif$^{R}$ | Spontaneous rifampicin-resistant mutant of R1818 | (Marie et al., 2017) |
| *coiA* | R1818 *coiA*::*aph* (Kan$^{R}$) | This study<br>R4526 |
| *coiA*$^{D95A}$ | R1818 *coiA* with D95A mutation | This study<br>R2533 |
| *coiA*$^{D105A}$ | R1818 *coiA* with D105A mutation | This study<br>R2534 |
| *radA* | R1818 *radA*::*aadA* ? (Spec$^{R}$) | (Marie et al., 2017)<br>R2194 |
| *coiA radA* | R1818 *coiA*::*aph radA*::*aadA* (Kan$^{R}$ Spec$^{R}$) | This study<br>R4527 |
| R304 | Used as template for PCRc and PCRe | R304 |
| | ***Streptococcus mitis*** | |
| NCTC 12261 | NCTC 12261 [NS 51] | (Carlsson, 1968) |
| rif$^{R}$ (NCTC 12261) | Spontaneous rifampicin-resistant mutant of NCTC 12261 | This study |
| | ***Legionella pneumophila*** | |
| Paris | Paris Outbreak isolate CIP107629. | CNR Lyon |
| Paris_S | Spontaneous streptomycin-resistant mutant of Paris<br>Used as template to amplify the Strep$^{R}$ *rpsL* tDNA in Figure 1B. | This study |
| WT (*Lp*) | Paris *rocC*$_{TAA}$<br>Constitutively competent Paris mutant with TAA stop codon between the eleventh and twelfth codons of *rocC*. Denoted as wild-type in this study. | (Juan et al., 2015) |
| *yraN* (*Lp*) | Paris *rocC*$_{TAA}$ *yraN*::*aph* (Kan$^{R}$) | This study |
| *comM* (*Lp*) | Paris *rocC*$_{TAA}$ *comM*::*aacC1* (Gent$^{R}$) | This study |
| *yraN comM* (*Lp*) | Paris *rocC*$_{TAA}$ *yraN*::*aph comM*::*aacC1* (Kan$^{R}$ Gent$^{R}$) | This study |
| *yraN +yraN* | Paris *rocC*$_{TAA}$ *yraN*::*aph* pLPP::*lacI*$^{Q}$-P*tac-yraN-flag-aacC1* (Kan$^{R}$ Gent$^{R}$)<br>Complementation clone of *yraN* (*Lp*) | This study |
| *yraN +yraN*$^{D37A}$ | Paris *rocC*$_{TAA}$ *yraN*::*aph* pLPP::*lacI*$^{Q}$-P*tac-yraN*$^{D37A}$*-flag-aacC1* (Kan$^{R}$ Gent$^{R}$)<br>Complementation clone of *yraN* (*Lp*) with D37A mutation | This study |
| *yraN +yraN*$^{E50A}$ | Paris *rocC*$_{TAA}$ *yraN*::*aph* pLPP:: *lacI*$^{Q}$- P*tac-yraN*$^{E50A}$*-flag-aacC1* (Kan$^{R}$ Gent$^{R}$)<br>Complementation clone of *yraN* (*Lp*) with E50A mutation | This study |
| Lp02 | Philadelphia-1 *rpsL hsdR thyA* mutant | (Berger and Isberg, 1993) |

| | | |
|---|---|---|
| Lp02 *yraN* | Lp02 *yraN*::*aph* (Kan$^R$) | This study |
| Lp02 *comM* | Lp02 *comM*::*aph* (Kan$^R$) | This study |
| Lp02 *comEC* | Lp02 *comEC*::*aph* (Kan$^R$) | This study |
| Lp02 *comM* | Lp02 *comM*::*aph* (Kan$^R$) | This study |
| Phil-1 p45 Gent$^R$ | Philadelphia Outbreak isolate (1976)<br>Carrying the p45 element with the *aacC1* gene (Gent$^R$) | (Fraser et al., 1977) |
| | ***Acinetobacter baylyi*** | |
| WT (*Ab*) | ADP1 strain, also known and labeled as BD413<br>Denoted as wild-type *Acinetobacter baylyi* in this study. | (Juni and Janik, 1969)<br>BD413 |
| rif$^R$ (*Ab*) | Spontaneous rifampicin-resistant mutant of BD413, *rpoB* S531F | This study |
| *yraN* (*Ab*) | BD413 *yraN*::*aacC4* (Apra$^R$) | This study |
| *comM* (*Ab*) | BD413 *comM*::*aacC1* (Gent$^R$) | This study |
| *comM yraN* (*Ab*) | BD413 *yraN*::*aacC4 comM*::*aacC1* (Apra$^R$ Gent$^R$) | This study |
| *yraN* +*yraN* (*Ab*) | BD413 *yraN*::*aacC4* pMMB207C *yraN* (Apra$^R$ Cm$^R$)<br>Complementation clone of *yraN* (*Ab*) | This study |
| *yraN* +*yraN*$^{E60A}$ | BD413 *yraN*::*aacC4* pMMB207C *yraN*$^{E60A}$ (Apra$^R$ Cm$^R$)<br>Complementation clone of *yraN* (*Ab*) with E60A mutation | This study |
| WT (*mutS Ab*) | BD413 *mutS*::*aph* (Kan$^R$)<br>Denoted as wild-type *Acinetobacter baylyi* in the recombination event mapping assay. | This study |
| *yraN* (*mutS Ab*) | BD413 *mutS*::*aph yraN*::*aacC4* (Kan$^R$ Apra$^R$) | This study |
| *comM* (*mutS Ab*) | BD413 *mutS*::*aph comM*::*aacC1* (Kan$^R$ Gent$^R$) | This study |
| *yraN comM* (*mutS Ab*) | BD413 *mutS*::*aph yraN*::*aacC4 comM*::*aacC1* (Kan$^R$ Apra$^R$ Gent$^R$) | This study |
| | ***Acinetobacter nosocomialis*** | |
| M2 | Wild-type *Acinetobacter nosocomialis* | (Niu et al., 2008) |
| M2 rif$^R$ | Spontaneous rifampicin-resistant mutant of M2, *rpoB* Q522L | (Godeux et al., 2022) |
| M2 *yraN* | M2 *yraN*::*aacC4* (Apra$^R$) | This study |
| M2 *comM* | M2 Δ*comM* | (Tuffet et al., 2024) |
| M2 *yraN comM* | M2 *yraN*::*aacC4* Δ*comM* (Apra$^R$) | This study |
| M2 *lacZ-tet* | M2 *attTn7*::*lacZ-tetA* (Tet$^R$) | This study |
| M2 *yraN lacZ-tet* | M2 *attTn7*::*lacZ-tetA yraN*::*aacC4* (Tet$^R$ Apra$^R$) | This study |
| M2 *comM lacZ-tet* | M2 *attTn7*::*lacZ-tetA* Δ*comM* (Tet$^R$) | This study |
| M2 *yraN comM lacZ-tet* | M2 *attTn7*::*lacZ-tetA yraN*::*aacC4* Δ*comM* (Tet$^R$ Apra$^R$) | This study |
| M2 *lacZ*$^{dlt1}$-*kan* | M2 *attTn7*::*lacZ*$_{TAA}$-*aph* (Kan$^R$)<br>The 7-bp deletion in *lacZ* is at 250 nt from the stop codon of *aph*.<br>Used to amplify tDNA for comigration marker experiment | This study |
| M2 *lacZ*$^{dlt2}$-*kan* | M2 *attTn7*::*lacZ*$_{TAA}$-*aph* (Kan$^R$)<br>The 7-bp deletion in *lacZ* is at 1600 nt from the stop codon of *aph*. | This study |

| | | |
|---|---|---|
| | Used to amplify tDNA for comigration marker experiment | |
| M2 *lacZ*$^{dlt3}$-*kan* | M2 *attTn7*::*lacZ*$_{TAA}$-*aph* (Kan$^R$)<br>The 7-bp deletion in *lacZ* is at 3000 nt from the stop codon of *aph*.<br>Used to amplify tDNA for comigration marker experiment | This study |
| | ***Vibrio cholerae*** | |
| WT (*Vc*) | E7946 Sm$^R$, Δ*lacZ*::*lacIq*, P$_{tac}$-*tfoX*, Δ*luxO*, *pilA*$^{S67C}$, ΔVC1807::Zeo$^R$<br>Denoted as wild-type *Vibio cholerae* in this study | (Chlebek et al., 2019) |
| *yraN* (*Vc*) | E7946 Sm$^R$, Δ*lacZ*::*lacIq*, P$_{tac}$-*tfoX*, Δ*luxO*, *pilA*$^{S67C}$, ΔVC1807::Zeo$^R$, Δ*yraN::*Tm$^R$ (*yraN* = VC0580) | This study<br>TND1026 |
| *yraN* +*yraN* (*Vc*) | E7946 Sm$^R$, Δ*lacZ*::*lacIq*, P$_{tac}$-*tfoX*, Δ*luxO*, *pilA*$^{S67C}$, ΔVC1807::Zeo$^R$, Δ*yraN::*Tm$^R$, igVCA0265-VCA0266::Carb$^R$-*araC*-P$_{araBAD}$-*yraN* | This study<br>TND5041 |
| *comM* (*Vc*) | E7946 Sm$^R$, Δ*lacZ*::*lacIq*, P$_{tac}$-*tfoX*, Δ*luxO*, *pilA*$^{S67C}$, ΔVC1807::Zeo$^R$, Δ*comM*::Carb$^R$ (*comM* = VC0032) | (Dalia and Dalia, 2025)<br>TND2245 |
| *yraN comM* (*Vc*) | E7946 Sm$^R$, Δ*lacZ*::*lacIq*, P$_{tac}$-*tfoX*, Δ*luxO*, *pilA*$^{S67C}$, ΔVC1807::Zeo$^R$, Δ*yraN::*Tm$^R$, Δ*comM*::Carb$^R$ | This study<br>TND2249 |
| rif$^R$ (*Vc*) | E7946 Sm$^R$, Δ*lacZ*::*lacIq*, P$_{tac}$-*tfoX*, Δ*luxO*, *pilA*$^{S67C}$, *rpoB*$^{Q513P}$ (Rif$^R$) | This study |
| ΔVC1807::Erm$^R$ | E7946 Sm$^R$, ΔVC1807::Erm$^R$ | This study |
| | ***Escherichia coli*** | |
| DH5α | F$^-$ *supE44* Δ*lac*U169 (Φ*lacZ*ΔM15) *recA*1 *endA*1 *hsdR*17 *thi-1 gyrA*96 *relA*1<br>Cloning strain | Lab collection |
| BTH101 | F$^-$ *cya$^-$99 araD*139 *galE*15 *galK*16 *hsdR*2 *mcrA*1 *mcrB*1 *rpsL*1 (Strep$^R$)<br>Test strain for bacterial two-hybrid assay | (Karimova et al., 1998) |
| YraN-T25 YraN-T18 | BTH101 pKNT25-YraN pUT18-YraN (Kan$^R$ Amp$^R$) | This study |
| YraN-T25 RecA-T18 | BTH101 pKNT25-YraN pUT18-RecA (Kan$^R$ Amp$^R$) | This study |
| YraN-T25 ComM-T18 | BTH101 pKNT25-YraN pUT18-ComM (Kan$^R$ Amp$^R$) | This study |
| RecA-T25 YraN-T18 | BTH101 pKNT25-RecA pUT18-YraN (Kan$^R$ Amp$^R$) | This study |
| RecA-T25 RecA-T18 | BTH101 pKNT25-RecA pUT18-RecA (Kan$^R$ Amp$^R$) | This study |
| RecA-T25 ComM-T18 | BTH101 pKNT25-RecA pUT18-ComM (Kan$^R$ Amp$^R$) | This study |
| BL21 Rosetta (DE3) | BL21-Rosetta(DE3)-pLysS (Cm$^R$) (Novagen) | Lab collection |
| BL21 pKHS-*yraN* | BL21 Rosetta (DE3) carrying the pKHS-*yraN* plasmid (Cm$^R$ Kan$^R$)<br>Used to produce YraN$_{Lp}$ | This study |
| BL21 pKHS-*yraN*$^{E50A}$ | BL21 Rosetta (DE3) carrying the pKHS-*yraN*$^{E50A}$ plasmid (Cm$^R$ Kan$^R$)<br>Used to produce YraN$_{Lp}$ with E50A mutation | This study |
| BL21 pKHS-*coiA* | BL21-Rosetta(DE3) carrying the pKHS-*coiA* plasmid (Cm$^R$ Kan$^R$)<br>Used to produce CoiA$_{Sp}$ | This study |
| BL21 pKHS-*coiA*$^{D95A}$ | BL21-Rosetta(DE3) carrying the pKHS-*coiA* plasmid (Cm$^R$ Kan$^R$) | This study |

| | | |
|---|---|---|
| | Used to produce $CoiA_{Sp}$ with D95A mutation | |
| BL21 pKHS-*coiA*$^{E105A}$ | BL21-Rosetta(DE3) carrying the pKHS-*coiA* plasmid ($Cm^R$ $Kan^R$)<br>Used to produce $CoiA_{Sp}$ with E105A mutation | This study |
| BL21 pKHS-*recA* | BL21 (DE3) carrying the pKHS-*recA* plasmid ($Cm^R$ $Kan^R$)<br>Used to produce RecA for the nuclease assays | (Marie et al., 2017) |
| BL21 pKHS-*dprA* | BL21 (DE3) carrying the pKHS-*dprA* plasmid ($Cm^R$ $Kan^R$)<br>Used to produce DprA for the nuclease assays | (Quevillon-Cheruel et al., 2012) |

**Table S4**. Plasmids used in this study

| Name | Properties | Reference |
|---|---|---|
| pMMB207C | *ori*$_{RSF1010}$ *cat IncQ lacI*$^{Q}$ *Ptac* | (Morales et al., 1991) |
| pMMB207C *yraN* | *ori*$_{RSF1010}$ *cat Ptac-yraN*$_{Ab}$*-flag* | This study |
| pMMB207C *yraN*$^{E60A}$ | *ori*$_{RSF1010}$ *cat Ptac-yraN*$_{Ab}^{E60A}$*-flag* | This study |
| pX5K | Derivative of the *ori*$_{RSF1010}$ pMMB207αb-Km-14 *mobA*::*aph* plasmid (Kan$^{R}$)<br>Used as tDNA in Figure S1A | (Segal and Shuman, 1998) |
| pLS1 | pMV158 derivative, rolling-circle type replicative plasmid (Tet$^{R}$)<br>Used as tDNA in Figure S1B | (Stassi et al., 1981) |
| pKNT25 | For N-terminal fusions to T25 fragment of CyaA | (Karimova et al., 1998) |
| pUT18 | For N-terminal fusions to T18 fragment of CyaA | (Karimova et al., 1998) |
| pKNT25-RecA | pKNT25 with N-terminal *recA*-T25 (Kan$^{R}$) | This study |
| pKNT25-YraN | pKNT25 with N-terminal *yraN*-T25 (Kan$^{R}$) | This study |
| pUT18-RecA | pUT18 with N-terminal *recA*-T18 (Amp$^{R}$) | This study |
| pUT18-YraN | pUT18 with N-terminal *yraN*-T18 (Amp$^{R}$) | This study |
| pUT18-ComM | pUT18 with N-terminal *comM*-T18 (Amp$^{R}$) | This study |
| pUC18 | *ori*$_{pUC18}$ *lacZ*α *bla* (Amp$^{R}$)<br>Used as substrate for nuclease assay in Figures 2A and 3D | (Norrander et al., 1983) |
| pKHS | pET28 derivative (Kan$^{R}$) | (Eckert et al., 2010) |
| pKHS-*yraN* | pKHS carrying the *yraN* ORF from *Lp* fused to a (C-ter) His6 tag (Kan$^{R}$) | This study |
| pKHS-*yraN*$^{E50A}$ | pKHS carrying the *yraN* ORF from *Lp* with E50A substitution fused to a (C-ter) His6 tag (Kan$^{R}$) | This study |
| pKHS-*coiA* | pKHS carrying the *coiA* ORF from *Sp* | This study |
| pKHS-*coiA*$^{D95A}$ | pKHS carrying the *coiA* ORF from *Sp* with D95A substitution | This study |
| pKHS-*coiA*$^{E105A}$ | pKHS carrying the *coiA* ORF from *Sp* with E105A substitution | This study |
| pKHS-*recA* | pKHS carrying the *recA* ORF for RecA protein production | (Marie et al., 2017) |
| pKHS-*dprA* | pKHS carrying the *dprA* ORF for DprA protein production | (Marie et al., 2017) |

**Table S5.** Oligonucleotides used in this study

| Name | Sequence | Use |
|---|---|---|
| LH13_lpp3065_P1 | CAACGCTTTTCAAGGTATTAAGAGC | Forward primer to amplify a 2kB fragment uptstream of *yraN* from *Lp* |
| LH14_lpp3065_P2-tail-pKD4 | GGAACTTCGAAGCAGCTCCAGCCTACA-CAATCGCCAATTGTTCAGCAAACTTTCC | Reverse primer to amplify a 2kB fragment uptstream of *yraN* from *Lp* |
| LH15_lpp3065_P3-tail-pKD4 | GAACTAAGGAGGATATTCA-TATGGACCATGGCGGTTAAAGAACGCAT-TTGATGCAGG | Forward primer to amplify a 2kB fragment downstream of *yraN* from *Lp* |
| LH16_lpp3065_P4 | GACAACATTGCAGATCTCATCCATG | Reverse primer to amplify a 2kB fragment downstream of *yraN* from *Lp* |
| LH45_recA_P1 | GCAATTAGTGACTGCGGAATC | Forward primer to amplify a 2kB fragment uptstream of *recA* from Lp |
| LH46_recA_P2 | GGAACTTCGAAGCAGCTCCAGCCTACA-CAATCCAATATCCCGTGATACCGTGC | Reverse primer to amplify a 2kB fragment uptstream of *recA* from *Lp* |
| LH47_recA_P3 | GAACTAAGGAGGATATTCA-TATGGACCATGGCGTTGTCTGTGTT-GGCTAGTTC | Forward primer to amplify a 2kB fragment downstream of *recA* from *Lp* |
| LH48_recA_P4 | CCTTTATCTCCTACCTGACCACC | Reverse primer to amplify a 2kB fragment downstream of *recA* from *Lp* |
| LH55_dprA_P1 | CTCCGGATTCAGCATCACC | Forward primer to amplify a 2kB fragment uptstream of *dprA* from *Lp* |
| LH56_dprA_P2 | GGAACTTCGAAGCAGCTCCAGCCTACA-CAATCGGGCGAGGAGGGGTTGCAG | Reverse primer to amplify a 2kB fragment uptstream of *dprA* from *Lp* |
| LH57_dprA_P3 | GAACTAAGGAGGATATTCA-TATGGACCATGGCCTACTTATT-GCAGCAGGGTGC | Forward primer to amplify a 2kB fragment downstream of *dprA* from *Lp* |
| LH58_dprA_P4 | CTACCACGTATTCGCGCTG | Reverse primer to amplify a 2kB fragment downstream of *dprA* from *Lp* |
| LH66_pUT18_F | AGCTCGAATTCAGCCGCCAGCG | Forward primer to amplify T18 from pUT18 plasmid by SLIC |
| LH67_pUT18_R | CATAGCTGTTTCCTGTGTGAAATTGT-TATC | Reverse primer to amplify T18 from pUT18 plasmid by SLIC |
| LH74_pKNT25_R | CATAGCTGTTTCCTGTGTG | Forward primer to amplify T25 from pUT25 plasmid by SLIC |
| LH75_pKNT25_F | AGCTCGAATTCAATGACCATGC | Reverse primer to amplify T25 from pUT25 plasmid by SLIC |
| LH76_yraN-pKNT25_F | CACA-CAGGAAACAGCTATGACCCAAGAAAAAG GAAAGTTTGCTG | Forward primer to amplify *yraN* for cloning in pKNT25 by SLIC |
| LH77_yraN-pKNT25_R | GCATGG-TCATTGAATTCGAGCTGCATCCTGCATC AAATGCGTTC | Reverse primer to amplify *yraN* for cloning in pKNT25 by SLIC |
| LH82_comM-pUT18_F | TCACA-CAGGAAACAGCTATGAGTCTCGCAT-TTACCAAAACGC | Forward primer to amplify *comM* for cloning in pUT18 by SLIC |

| | | |
|---|---|---|
| LH83_comM-pUT18_R | CTGGCGGCTGAATTCGAGCTTTTCGG-TAAGTGTAAATTTTGTTTG | Reverse primer to amplify *comM* for cloning in pUT18 by SLIC |
| LH98_recA-pUT18_F | TCACACAGGAAACAGCTATGGAAGA-GAATAAACAAAAAGCAC | Forward primer to amplify *recA* for cloning in pUT18 by SLIC |
| LH99_recA-pUT18_R | CTGGCGGCT-GAATTCGAGCTGTCATCTA-TAGTCTCAAAAAGATCTTC | Reverse primer to amplify *recA* for cloning in pUT18 by SLIC |
| LH102_recA-pKNT25_F | CACACAGGAAACAGCTATGGAAGAGAA-TAAACAAAAAGCAC | Forward primer to amplify *recA* for cloning in pKNT25 by SLIC |
| LH103_recA-pKNT25_R | GCATGG-TCATTGAATTCGAGCTGTCATCTA-TAGTCTCAAAAAGATCTTC | Reverse primer to amplify *recA* for cloning in pKNT25 by SLIC |
| LH118_YraN_D37A_F | ATCATTGCCGTTTAGGA-GAAATTGCCTTAATTATGCGTGAAG-GATCTTATC | Forward primer to replace codon coding for D37 to A37 in *yraN* |
| LH119_YraN_D37A_R | GATAAGATCCTTCACGCATAAT-TAAGGCAATTTCTCCTAAACGGCAATGAT | Reverse primer to replace codon coding for D37 to A37 in *yraN* |
| LH120_yraN_E50A_F | GTGAAGGATCTTATCTGGTTTTTATT-GCCGTTCGTTCGCGTTCAAATATG | Forward primer to replace codon coding for E50 to A50 in *yraN* |
| LH121_yraN_E50A_R | CATATTTGAACGCGAACGAACGGCAA-TAAAAACCAGATAAGATCCTTCAC | Reverse primer to replace codon coding for E50 to A50 in *yraN* |
| LH140_plpp0110_P1 | GCCAAACGCTATCAGGAGTTACCTC | P1 primer to clone *yraN* in pLPP |
| LH141_plpp0110_P2-tail-LacIq | CTTGCTGGCGTTCGGGAGCA-GAAGAGCATAGATTGTCCCATCTCCTT-GAACC | P2 primer to clone *yraN* in pLPP |
| LH142_plpp0110_P3-tail-CmR | GTTCCGCGTCCTTGCAATACTGTGTT-TACGCGAAGAATCAAGAATAACGCCAC | P3 primer to clone *yraN* in pLPP |
| LH143_plpp0110_P4 | CTGATGAGGCGCTGCGAAGAG | P4 primer to clone *yraN* in pLPP |
| LH144_plpp3065-La-cIq_F | TATGCTCTTCTGCTCCCGAACGCCAGCA AG | Forward primer to clone *yraN* in pLPP |
| LH145_plpp3065-CmR_R | GTAAACACAGTATTGCAAG-GACGCGGAAC | Reverse primer to clone *yraN* in pLPP |
| LH146_2kB-AttTn7_P1 | GTTCTTTCTGCTGTGCGAAAGGAC | P1 primer to insert *lacZ-atbR* cas-sette in *attTn7* from M2. Also used as forward primer to amplify tDNA for the comigration marker experiment |
| LH147_AttTn7-tail-lacz_P2 | ATTATACC-TAGGACTGAGCTAGCTGTCAACGT-GAAAGAAAATCTAGTTTGATAGTAGAG | P2 primer to insert *lacZ-atbR* cas-sette in *attTn7* from M2 |
| LH148_attTn7-tail-kan_P3-v4 | CATCAACCATATCAGCAAAAGT-GATACGGTAAAAGCGCTTGTG-TAGCGATATGAATG | P3 primer to insert *lacZ-atbR* cas-sette in *attTn7* from M2 |
| LH149_attTn7-2kB_P4 | CGTGCTCAACAAATTTCTAAGGC | P4 primer to insert *lacZ-atbR* cas-sette in *attTn7* from M2. Also used as reverse primer to amplify tDNA for the comigration marker experiment |
| LH150_TcR-tail-lacZ_F | CATTCATATCGCTACA-CAAGCGCTTTTAACATCGCAAAAGGTG-TAAATATTATAATGC | Forward primer to insert *lacZ-atbR* cassette in *attTn7* from M2 |

| | | |
|---|---|---|
| LH151_TcR-tail-attTn7_R | GCCAGTGTTACAACCAAT-TAACCAATTCT-GATACTTCCATTCAGGTCGAGGTGGCCCGGC | Reverse primer to insert *lacZ-atbR* cassette in *attTn7* from M2 |
| LH152_2k-TcR_P1 | TTCGCATTATCCGAACCATCCGC | P1 primer to amplify *tetA* cassette |
| LH153_2kb-TcR_P2 | CAGAATTGGTTAATTGGTT-GTAACACTGGC | P2 primer to amplify *tetA* cassette |
| LH154_TcR-2kB_P3 | TAAAAGCGCTTGTGTAGCGATATGAATG | P3 primer to amplify *tetA* cassette |
| LH155_strg-PRM_lacZ_F | TTGACAGCTAGCTCAGTCC-TAGGTATAATTTCACACAGGAAACA-GAATTCTTTAAGAAGGA-GATATACATATGGTCGTTTTACAA | F primer to add a strong promoter to *lacZ* |
| LH156_kan_R-V4 | CCGTATCACTTTTGCTGATATGGTTGATG | Reverse primer to amplify *aph* cassette |
| LH159_lacZ-dlt1_P2 | AACTGTTGGGAAGGGCGATC | Reverse primer to delete 7 bp at the beginning of *lacZ* gene |
| LH160_lacZ-dlt1-tailed_P3 | CCCGCACCGATCGCCCTTCCCAACAGT TGTGAATGGCGAATGGCGCTTTG | Forward primer to delete 7 bp at the beginning of *lacZ* gene |
| LH161_lacZ-dlt2_P2 | GCCTTCTTCCGCGTGCAGCAG | Reverse primer to delete 7 bp at the end of *lacZ* gene |
| LH162_lacZ-dlt2-tailed_P3 | CATCGCCATCTGCTGCACGCGGAAGAA GGCTGAATATCGACGGTTTCCATATGGG | Forward primer to delete 7 bp at the end of lacZ gene |
| LH163_lacZ-dlt3_P2 | CGTCTCTCCAGGTAGCGAAAG | Reverse primer to delete 7 bp at the middle of *lacZ* gene |
| LH164_lacZ-dlt3-tailed_P3 | AAAAAATGGCTTTCGCTACCTGGAGA-GACGTGATCCTTTGCGAATACGCC | Forward primer to delete 7 bp at the middle of *lacZ* gene |
| LH222_yraN-bay-lyi_P1 | CAGCCTTAACCGAGAGGTC | Forward primer to amplify a 1kB fragment uptstream of *yraN* from BD413 |
| LH223_yraN-baylyi_P2-tail-apra | TGCGGGAGGGCAAGGGCTCCAAG-GATCGGCATAGCTTCCTAGAGTTCGTGC | Reverse primer to amplify a 1kB fragment uptstream of *yraN* from BD413 |
| LH224_yraN-baylyi_P3-tail-apra | CCGCTCGCCAGTCGATTGGCT-GAGCTCATGACCGCTGCCAAT-GAGGTAC | Forward primer to amplify a 1kB fragment uptstream of *yraN* from BD413 |
| LH225_yraN-bay-lyi_P4 | GGAGGAAGAAATTAGCGACGC | Reverse primer to amplify a 1kB fragment uptstream of *yraN* from BD413 |
| LH228_comM-bay-lyi_P1 | CCCAATTGGCACAGCATTTG | Forward primer to amplify a 1kB fragment uptstream of *comM* from BD413 |
| LH229_comM-baylyi_P2-tl-gent | CGGTAAATTGTCA-CAACGCCGCCAGGTGGCCGCT-TACCCCTCAAGAGAATC | Reverse primer to amplify a 1kB fragment uptstream of *comM* from BD413 |
| LH230_comM-baylyi_P3-tl-gent | TCCGTTTCCACGG-TGTGCGTCCATGGGCAAGTGGCAAAA-TACTCGGTAAGCG | Forward primer to amplify a 1kB fragment downstream of *comM* from BD413 |
| LH231_comM-bay-lyi_P4 | GCAGCAGCGACAAAGCTATG | Reverse primer to amplify a 1kB fragment downstream of *comM* from |

| | | |
|---|---|---|
| | | BD413 |
| LH353_muts_ADP1_P1 | CCACGTCAAACCAATCAGTCG | Forward primer to amplify a 1kB fragment downstream of *mutS* from BD413 |
| LH364_mutS_BD413_P2 | CATCAACCATATCAGCAAAAGT-GATACGGGATAACTGCCAAGCAATGCG | Reverse primer to amplify a 1kB fragment downstream of *mutS* from BD413 |
| LH363_mutS_BD413_P3 | TTTCATTTGATGCTCGAT-GAGTTTTTCTAAGCA-TAAAGTCCAGCATGGCC | Forward primer to amplify a 1kB fragment uptstream of *mutS* from BD413 |
| LH356_mutS_ADP1_P4 | GGTCATGGCGACAGCTACC | Reverse primer to amplify a 1kB fragment uptstream of *mutS* from BD413 |
| LH232_gentR_F | TTGCCCATGGACGCACACCGTG | Forward primer to amplify *aacC1* cassette |
| LH233_GentR_R | GCCACCTGGCGGCGTTGTG | Reverse primer to amplify *aacC1* cassette |
| BBC2172 | TACAATGCATTCTTTTTCAACTGG | Amplification of the upstream region of *yraN* from *Vc* |
| BBC2173 | GTCGACGGATCCCCGGAATCCTAGAAT-TAACGAACACCATGC | |
| BBC2174 | GAAGCAGCTCCAGCCTACAAACGC-TATTACTGAAGGATAAAC | Amplification of the downstream region of *yraN* from *Vc* |
| BBC2175 | AATGATCTTGGCATCCCAGC | |
| BBC3267 | CAATTTCACACAGGATCCCGGGAG-GAGGTGTATTGATGAACAATGTGTGG-TTC | Amplification of *yraN* from *Vc* for complementation at ectopic site |
| BBC3268 | TGTAGGCTGGAGCTGCTTCT-TATCCTTCAGTAATAGCGTTTTTTAACC | |
| TMN0347 | GTTGAAGAGCAAACTGAATTCAACG | Amplification of the *rpoB* cassette from the rifR *Vc* strain |
| TMN0346 | GGTCGAGGTACTCTTCCTCAG | |
| BBC717 | AAATAGATTTGGTGACTTTACCTCC | Amplification of the erythromycin cassette from ∆VC1807::ErmR *Vibrio cholerae* strain |
| BBC718 | CTTTACGCCTGATTGTCTACAC | |
| MB137 | CGTCTAGGACACGCATGTCAAGA | Amplification of PCRc *rpoB (*4195 bp) (Marie et al., 2017) |
| MB138 | GGCGGTAGACGGATTTGAACC | |
| MB139 | TTCTTCTTGAATGGAGGGTTGAGC | Amplification of PCRe *rpoB* (3912 bp) (Marie et al., 2017) |
| MB140 | GTCCGTGAACGTATGTCTGTTCAGG | |
| rpsL_Fw | GCAGCTCCAGATGGCTCAATC | Amplification of a 4 kb fragment encompassing a Strep$^R$ *rpsL* allele used as tDNA in *Lp* |
| rpsL_Rv | CAACCATACATGTCCATATTGACCAC | |
| lpp2553_P1 | GTGTCGCCACAGCACCAAGATC | Construction of the *lpp2553*::*aacC4* mutant in *Lp* and amplification of the *lpp2553*::*aacC4* fragment used as Apra$^R$ tDNA in *Lp* |
| lpp2553_P4 | GCCCGCAAGCTACAAATACCATAG | |
| lpp2553_P2 | GGAACTTCGAAGCAGCTCCAGCCTACACAATCCGATATCAGAGAGGCTATGTACTCC | Construction of the *lpp2553*::*aacC4* mutant in *Lp* |

| | | |
|---|---|---|
| lpp2553_P3 | GAACTAAGGAGGATATTCATATGGACCATGGCAAGCCAACCTTGATGGAGAGC | |
| LH280_rpoB_M2/ADP1_12kB_R | GCTGTTACAGGCAAGATTTCAACAG | Amplification of a 11 kb fragment encompassing a *rpoB* rifR allele as tDNA in *Ab* |
| LH281_rpoB_M2/ADP1_10kB_F | GTTGGGGGTTCGAGTCCCTC | |
| CR63_rpoB_6kb_exc_F | TTCTGGCAATGCTGCTTTAG | Amplification of 6.4 kb rifR *rpoB-rpoC* fragment in M2 donor and BD413 recipient |
| CR64_rpoB_6kb_exc_R | TCACCACCTTGAATGAATGG | |
| CR66_rpoB_verif_CTG_M2 | GGTTGTTTTGRTCCATRAACA | Diagnosis PCR to differenciate BD413 *mutS* transformants and spontaneous rifR mutants |
| CR31_rpoB_6kb_F | CGTATGAAGTTCAACCGTCG | |
| MB530 | TTCCTAGACCATGGTCTGAAGGAAGTG | Amplification of 6.4 kb rifR *rpoB-rpoC* fragment in *S. mitis* donor |
| MB533 | GGGCTTCAAAGATTTCTTGGACACGAGG | |
| MB138 | GGCGGTAGACGGATTTGAACC | Amplification of 7.8 kb rifR *rpoB-rpoC* fragment in *S. pneumoniae* recipient |
| MB519 | TTATTCTACTGTTTCTTCCACAGTTTCAACG | |
| LH59_yraN_ABUW2846_P1 | CCGTCCTTGTGTACTCACAC | Primers used to construct the *yraN*::*aacC4* allele that was transformed into the M2 strain. |
| LH60_yraN_ABUW2846_P2 | GGAACTTCGAAGCAGCTCCAGCCTACACAATCGATAATTACTGGCGACCCATTCA | |
| LH61_yraN_ABUW2846-P4 | CTGCAACTCGTTCCAATTC | |
| LH62_yraN_ABUW2846-P3 | AACTAAGGAGGATATTCATATGGACCATGGCTTACAGCGCTACCCATCTTATC | |
| coiA1-Fw | TACAATACCAATCCTGTTCATTCAG | Forward primer to amplify a 2kB fragment uptstream of *coiA* from *Sp* |
| coiA2-Rev | CTGCTTTTCGCTAATCTCCATAA | Reverse primer to amplify a 2kB fragment uptstream of *coiA* from *Sp* |
| coiA_EagI-Fw | TTTTTCGGCCGATGTTTGTTGCGAGAGATGCTAGGGG | amplification of *coiA* ORF from *Sp*, Cloned after EagI+NotI digestion in pKHS digested by NotI |
| coiA_NotI-Rev | TTTTTGCGGCCGCTTATTCTACCATATTTTTCAAG | |
| coiA_D95A | GAACTTAAACAGATTGCGGCCGTATTTGTAAATGGCAATC | *coiA* directed mutagenesis |
| coiA_E105A | GGCAATCTAGCTCTAGCCGTTCAGTGTAGTCCCTTGC | *coiA* directed mutagenesis |
| yraN_Lp-IF-Fw | GGAGATATATATATGACCCAAGAAAAAGGAAAGTTTGCTG | Amplification of *yraN* ORF from *Lp*, cloned in pKHS vector by InFusion strategy |
| yraN_Lp-IF-Rev | GTGGTGGTGGTGGTGGCATCCTGCATCAAATGCGTTCTTTAACC | |
| pKHS-IF-Rev | CATATATATATCTCCTTCTTAAATTAATCGGCCGCAAGCTCTAG | Amplification of pKHS vector to clone *yraN* ORF by InFusion strategy |
| pKHS-IF-Fw | CACCACCACCACCACTGAGATCCGGCTGCTAAC | |
| yraN_E50A | CTTATCTGGTTTTTATTGCGGTTCGTTCGCGTTC | *yraN* directed mutagenesis |

| | | |
|---|---|---|
| Ovio1 | CTAGGGTCGGATCCTCTAGACAGCTCC ATGATCACTGGCACTGGTAGAATTCGG CCCATTAGCAAGGCCGGAAACGTCACC CTCCAGTTTCTCGCCTCTG | 100 mer oligonucleotide used as ssDNA, dsDNA (with Ovio7) or hybrid ss-dsDNA (with Ovio3, Ovio5) |
| Ovio3 | CAGAGGCGAGAAACTGGAGGGTGACGT TTCCGGCCTTGCTAATGGGCCGAATTCT ACCAGTGCCAGTGAT | 70 mer oligonucleotide used as hybrid ss-dsDNA with Ovio1 and Ovio14 |
| Ovio4 | TAGCAATGTAATCGTCTATGACGTTAAA CCATCGATAGCAGCACCGTAATCAGTA GCGACAGAATCAAGT | 70 mer oligonucleotide used as hybrid ss-dsDNA with Ovio50 |
| Ovio5 | CATGGAGCTGTCTAGAGGATCCGACCC TAG | 30 mer oligonucleotide used as hybrid ss-dsDNA with Ovio1 |
| Ovio7 | CAGAGGCGAGAAACTGGAGGGTGACGT TTCCGGCCTTGCTAATGGGCCGAATTCT ACCAGTGCCAGTGATCATGGAGCTGTC TAGAGGATCCGACCCTAG | 100 mer oligonucleotide used as dsDNA (with Ovio1) |
| Ovio9-Cy3 | AATCATGGTCATAGCTGTTTCCTGTGTG AATAGCAATGTAATCGTCTATGACGTTA AACCATCGATAGCAGCACCGTAATCAGT AGCGACAGAATCAAGT | 100 mer oligonucleotide with a Cy3 modification in 3', used as ssDNA |
| Ovio14 | TTCACACAGGAAACAGCTATGACCATGA TTCTGCATAGATCTAGGGTCGGATCCTC TAGACAGCTCCATGATCACTGGCACTG GTAGAATTCGGCCCATT | 100 mer oligonucleotide used as ssDNA or hybrid ss-dsDNA (with Ovio3, Ovio18, Ovio19 and Ovio50) |
| Ovio18 | AATGGGCCGAATTCTACCAGTGCCAGT GAT | 30 mer oligonucleotide used as hybrid ss-dsDNA with Ovio14 |
| Ovio19 | AATGGGCCGAATTCTACCAGTGCCAGT GATACTTGATTCTGTCGCTACTGATTAC GGTGC | 60 mer oligonucleotide used as hybrid ss-dsDNA (with Ovio14 and Ovio49) |
| Ovio46 | GCACCGTAATCAGTAGCGACAGAATCA AGT | 30 mer oligonucleotide used as hybrid ss-dsDNA with Ovio19 |
| Ovio50 | AATGGGCCGAATTCTACCAGTGCCAGT GATACTTGATTCTGTCGCTACTGATTAC GGTGCTGCTATAATCATGGTCATAGCTG TTTCCTGTGTGAA | 96 mer oligonucleotide used as hybrid ss-dsDNA with Ovio14 and Ovio4 |
| Cy3-Ovio51 | TGCTTCCGGCTCGTATGTTGTGTGGAAT TGTGAGCGGATAACAATTTCACACAGGA AACAGCTATGACCATGATTACGAATTCG AGCTCGGTACCCGGGG | 100 mer oligonucleotide with a Cy3 modification in 5', used to form a D-Loop with pUC18 |
| Cy3-cc13up | GACTGCATCACATTTGCATCATACATGC ATTATTTCCCTCTTGCAAGCCCGGTTTC CGTCCGTTTTAGCTCATTTTCTGCATCA TTGTAGCACCATCATAGCATTATAG | 100 mer oligonucleotide with a Cy3 modification in 5', used as a negative control for the D-Loop experiment |
| M13mp/pUC (-40) Primer | GTTTTCCCAGTCACGAC | commercial primer used for the primer extension experiment |
| Ovio25 | CTAGACAGCTCCATGATCACTGGCACT GG | 29 mer primer used as ss-dsDNA with Ovio26 |
| Ovio26 | CCAGTGCCAGTGAT | 14 mer primer used as ss-dsDNA with Ovio25, Ovio29, Ovio38 and Ovio39 |
| Ovio27 | CTAGACAGCTCCATGTACACTGGCACT | 29 mer primer used as ss-dsDNA |

| | | |
|---|---|---|
| | GG | with Ovio28 |
| Ovio28 | CCAGTGCCAGTGTA | 14 mer primer used as ss-dsDNA with Ovio27 |
| Ovio29 | CTAGACAGCTCCATCATCACTGGCACTG G | 29 mer primer used as ss-dsDNA with Ovio26 |
| Ovio30 | CTAGACAGCTCCATCATGACTGGCACT GG | 29 mer primer used as ss-dsDNA with Ovio31 |
| Ovio31 | CCAGTGCCAGTCAT | 14 mer primer used as ss-dsDNA with Ovio30 |
| Ovio32 | CTAGACAGCTCCATGATTACTGGCACTG G | 29 mer primer used as ss-dsDNA with Ovio33 |
| Ovio33 | CCAGTGCCAGTAAT | 14 mer primer used as ss-dsDNA with Ovio32 |
| Ovio34 | CTAGACAGCTCCATGAACACTGGCACT GG | 29 mer primer used as ss-dsDNA with Ovio35 |
| Ovio35 | CCAGTGCCAGTGTT | 14 mer primer used as ss-dsDNA with Ovio34 |
| Ovio36 | CTAGACAGCTCCATGTTCACTGGCACTG G | 29 mer primer used as ss-dsDNA with Ovio37 |
| Ovio37 | CCAGTGCCAGTGAA | 14 mer primer used as ss-dsDNA with Ovio36 |
| Ovio38 | CTAGACAGCTCCAGGATCACTGGCACT GG | 29 mer primer used as ss-dsDNA with Ovio26 |
| Ovio39 | CTAGACAGCTCCTAGATCACTGGCACT GG | 29 mer primer used as ss-dsDNA with Ovio26 and Ovio40 |
| Ovio40 | TAGGAGCTGTCTAG | 14 mer primer used as ss-dsDNA with Ovio39 |
